# CryoMDM; Molecular simulation Driven structural Matching Approach to Estimate Variety of Atomic Three-dimensional Model Based on Noisy cryoEM Single Particle Images

**DOI:** 10.1101/2024.10.31.621264

**Authors:** Atsushi Tokuhisa, Yoshinobu Akinaga, Yoko Sasakura, Kei Terayama, Shigeyuki Matsumoto, Takayuki Kato, Yasushi Okuno

## Abstract

Understanding the structure of biomolecules is essential for comprehending life phenomena. In recent years, cryo-electron microscopy (cryoEM) single-particle structure analysis has made remarkable advancements. The single-particle images obtained from cryoEM measurements have a poor signal-to-noise ratio, and existing methods rely on multi-image analysis, which averages multiple measurement data to achieve high-resolution structures. However, this multi-image approach statistically processes several measurement images, causing the unique structural features of individual data to be lost. To overcome this inherent challenge, we developed a molecular simulation-driven structural matching method, cryoMDM, which identifies the plausible structure for cryoEM two-dimensional images from a vast number of three-dimensional (3D) structural model candidates generated through molecular simulations. The innovative aspect of this method is that it enables the estimation of 3D atomic models from a single-particle image, going beyond merely generating a 3D electron density map. By linking each cryoEM particle image to a 3D structure in the simulation space, we can directly connect cryoEM measurement data to continuous structural changes in biomolecules. Applying this method to the spike protein derived from SARS-CoV-2, we successfully captured various intermediate structures of the spike protein, revealing critical information about the early stages of viral entry.

## Introduction

Various biological phenomena are supported by numerous biomolecules functioning both inside and outside of cells[1,2]. For biomolecules to exhibit their diverse functions, the flexibility to adopt multiple conformational states is crucial[3,4,5]. Therefore, understanding the diverse three-dimensional (3D) structural states of biomolecules is essential for comprehending life phenomena[6, 7].Cryo-electron microscopy (cryoEM) has made remarkable advancements in recent years as a powerful structural analysis technique[8, 9, 10]. In cryoEM single-particle structure analysis, two-dimensional (2D) projection images of biomolecular structures are obtained by irradiating electron beams on ice-embedded samples that have been rapidly frozen from solution[11]. The captured images (micrographs) reflect the dynamic properties of biomolecules, showing single-particle images from various orientations. This indicates that cryoEM has a very high potential to experimentally capture the diverse conformational states of biomolecules [12, 13].

Biological samples are susceptible to damage from strong electron beams, resulting in a low signal-to-noise (S/N) ratio for the micrographs obtained through cryoEM. In single-particle structure analysis, a representative method for determining the three-dimensional structures of biomolecules using cryoEM, this low S/N ratio is overcome through multi-image analysis [14]. This approach improves the S/N ratio by using a vast number of single-particle images picked from micrographs to generate high-resolution 3D density maps [15]. Recently, AI technologies have also been reported that automate these data analysis steps to create 3D density maps from 2D images [16, 17, 18].

However, these multi-image analysis approaches statistically process multiple measurement images, which means that while they can enhance the resolution of common structural features across images, the unique diverse structural states of individual measurements are averaged out, resulting in a loss of information [19, 20]. In other words, the structures obtained may be biased towards those that are more prevalent under the sample preparation conditions, while flexible structural components might remain invisible. This issue persists even when using image-generating AI, as any structural information not included in the training data cannot be estimated, leaving it as an inherent challenge.

To overcome this inherent challenge, we developed a molecular simulation-driven structural matching method called cryoMDM, which identifies structures that closely match the observed 2D single-particle images from cryoEM using a vast number of 3D structural model candidates generated through molecular dynamics (MD) simulations [21,22]. This method achieves single-image analysis by estimating a 3D atomic model from a single cryoEM particle image for the first time.

Our approach involves creating virtually all-directional 2D projection images from the diverse 3D structures sampled through MD simulations. We then perform comprehensive image matching with cryoEM measurement data, enhanced in quality using Variational Autoencoders (VAE) [23], to link individual experimental single-particle images to their corresponding three-dimensional structures. Furthermore, this method connects each cryoEM image to a 3D structure in a simulation space that incorporates time information, allowing us to directly relate cryoEM measurement data to the continuous structural changes of biomolecules.

As a result, we can capture intermediate states of biomolecules that exist between the stable structural states determined by conventional single-particle structure analysis, providing a comprehensive understanding of the behavior of biomolecules in solution.

In this study, we demonstrated the usefulness of the proposed method using the spike protein derived from SARS-CoV-2 [24]The spike protein is involved in the early stages of viral entry into host cells, and it has been experimentally shown to exist in two stable structural states: the open state (Open) and the closed state (Close), with the receptor-binding domain (RBD) responsible for adhesion to the host cell [25]. By applying the cryoMDM method to the experimental single-particle images of the spike protein, we successfully captured a variety of intermediate structures that are difficult to observe under normal experimental conditions, thereby revealing the complete transition between the two stable structural states.

## Result

### Overview of Simulation-Driven Structural Matching Method: cryoMDM

CryoMDM is an entirely new single-image dynamic structure analysis method that directly estimates a 3D atomic model from an individual 2D single-particle image obtained through cryoEM. We approached this problem as one of linking individual 2D single-particle images acquired by cryoEM to the spatiotemporal scale of a 3D atomic model, aiming to develop a corresponding analysis method. In particular, to achieve this, we must technically overcome issues related to the noise in the 2D single-particle images from cryoEM measurements, the generation of 3D structural models, and the matching of 2D single-particle images with 3D structural models. In this study, we implemented dynamic structure analysis from a single image by exploring and identifying the most similar structures among diverse 2D projection images of 3D structures generated by MD, using FT-image matching on cryoEM measured 2D single-particle images that underwent quality enhancement via VAE. Specifically, we linked cryoEM single-particle images obtained from experiments with 3D structures generated through MD simulations based on a four-step analysis workflow shown in Figure 1. Below, we outline the detailed procedures of this method.

**Figure 1.**
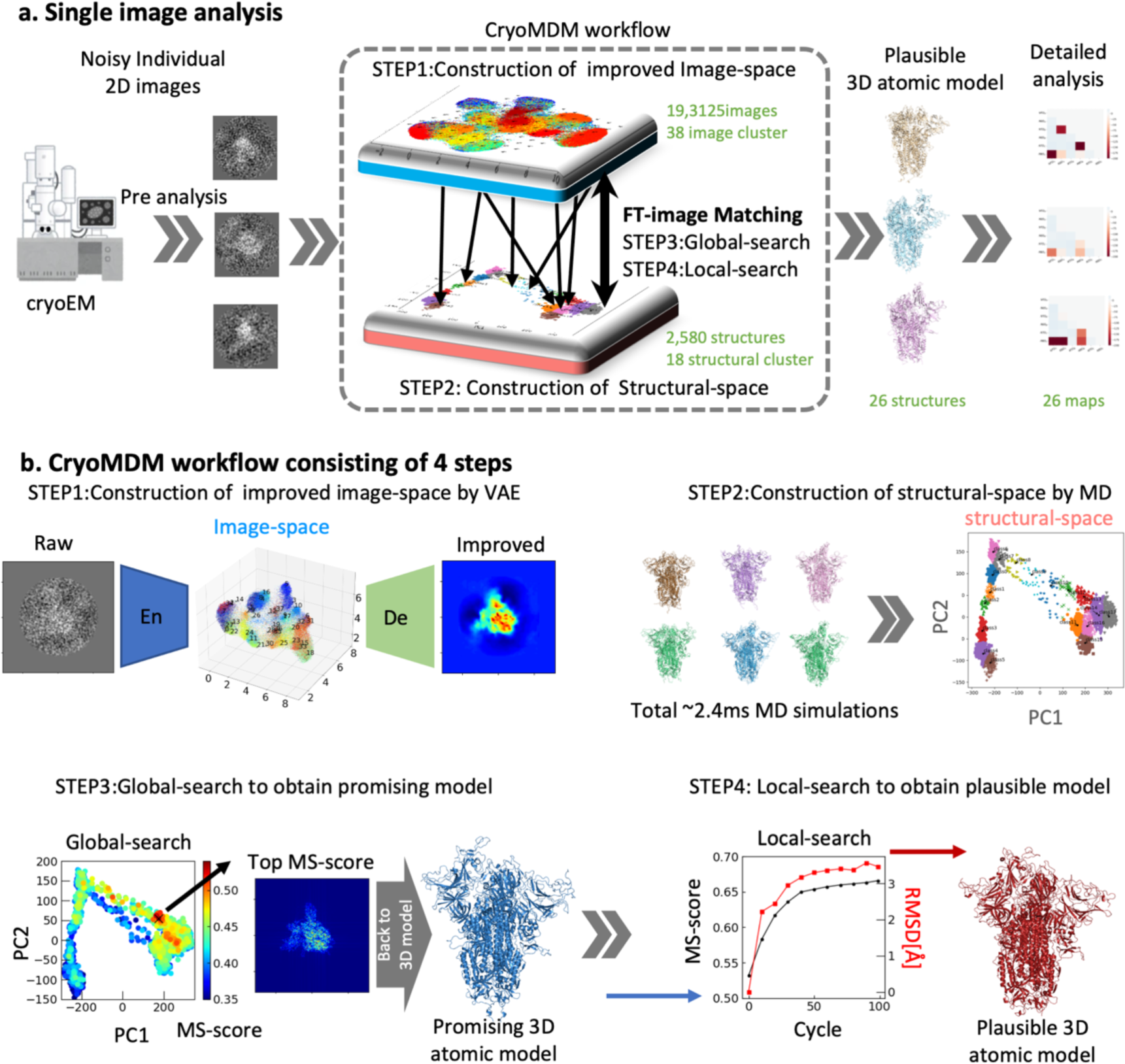
(a) Overview of cryoMDM single image analysis. (b) Conceptual diagram of the cryoMDM workflow consisting of 4 steps. In STEP 1, an improved image space is constructed. In STEP 2, a structural space composed of diverse structures is created using MD. The image space and the structural space are linked through FT-image Matching. At this stage, for efficiency, a global search is performed in STEP 3, followed by a local search in STEP 4.

The individual single-particle images (2D images) obtained from cryoEM measurements are noisy, making it difficult to map them directly to 3D structures. Therefore, in Step 1, we improve the quality of the individual single-particle images by training the cryoEM 2D images using a Variational Autoencoder (VAE) [23]. To capture large-scale structural changes according to the manifold hypothesis, we select representative images through image clustering in the VAE latent space (image space), which will serve as the target for subsequent calculations.

Next, in Step 2, we construct the structural space using molecular dynamics (MD) simulations[21]. The atomic-level 3D structures possess a very high degree of freedom, resulting in an extensive spatiotemporal scale for the 3D structural space. While MD is a powerful method for capturing 3D atomic models in spatiotemporal scales, its computational cost is very high, leading to a narrow sampling of the structural space with standard MD alone. Therefore, in this implementation, we combine standard MD with targeted MD[26] to build a broader structural space.

Next, to connect the constructed image space and structural space, we comprehensively calculate the similarity scores between the improved experimental 2D images and a set of 2D projection images generated from 3D atomic structures created through MD simulations (with 10,240 projections over 180 degrees). The 3D atomic model that shows the highest similarity score is selected as the promising structure (Step 3: global search). It is important to emphasize that we use a uniquely designed similarity score based on FT-image analysis for this similarity assessment. This score effectively suppresses the degrees of freedom related to translation and rotation while being robust to noise. In Step 4, we take the promising structure obtained from the global search as the initial structure and explore structures in its vicinity that better match the target single-particle image to obtain the final model structure (Step4: local search). During this local search, we apply a Modified PaCS-MD algorithm with the similarity score from FT-image matching as a constraint, allowing for efficient search of structural states that demonstrate even higher similarity scores. Given that both the MD simulations and comprehensive matching require significant computational resources, we utilized the cutting-edge the supercomputer Fugaku[27] for the demonstration of the cryoMDM workflow shown in the following sections.

### Evaluation of Single-Image Analysis Flow Using Synthetic Data

In this method, it is crucial to estimate the correct 3D structure from an individual single-particle image with the correct molecular orientation based on the similarity of FT-image matching. Therefore, we first conducted a validation of FT-image matching using synthetic data as the target image.

First, we prepared a set of 2,580 diverse structures that span the structural space including both Open and Close states of the Spike protein, using publicly available MD simulation trajectories [28, 29, 30] (PCA plot). We applied the X-mean clustering method to these structures, resulting in 18 representative structures (Figure 2a). This set of representative structures included typical configurations traversing between the Close state (upper part of the PC1-PC2 plane), intermediate states (lower left part of the PC1-PC2 plane), and the Open state (lower right part of the PC1-PC2 plane).

**Figure 2.**
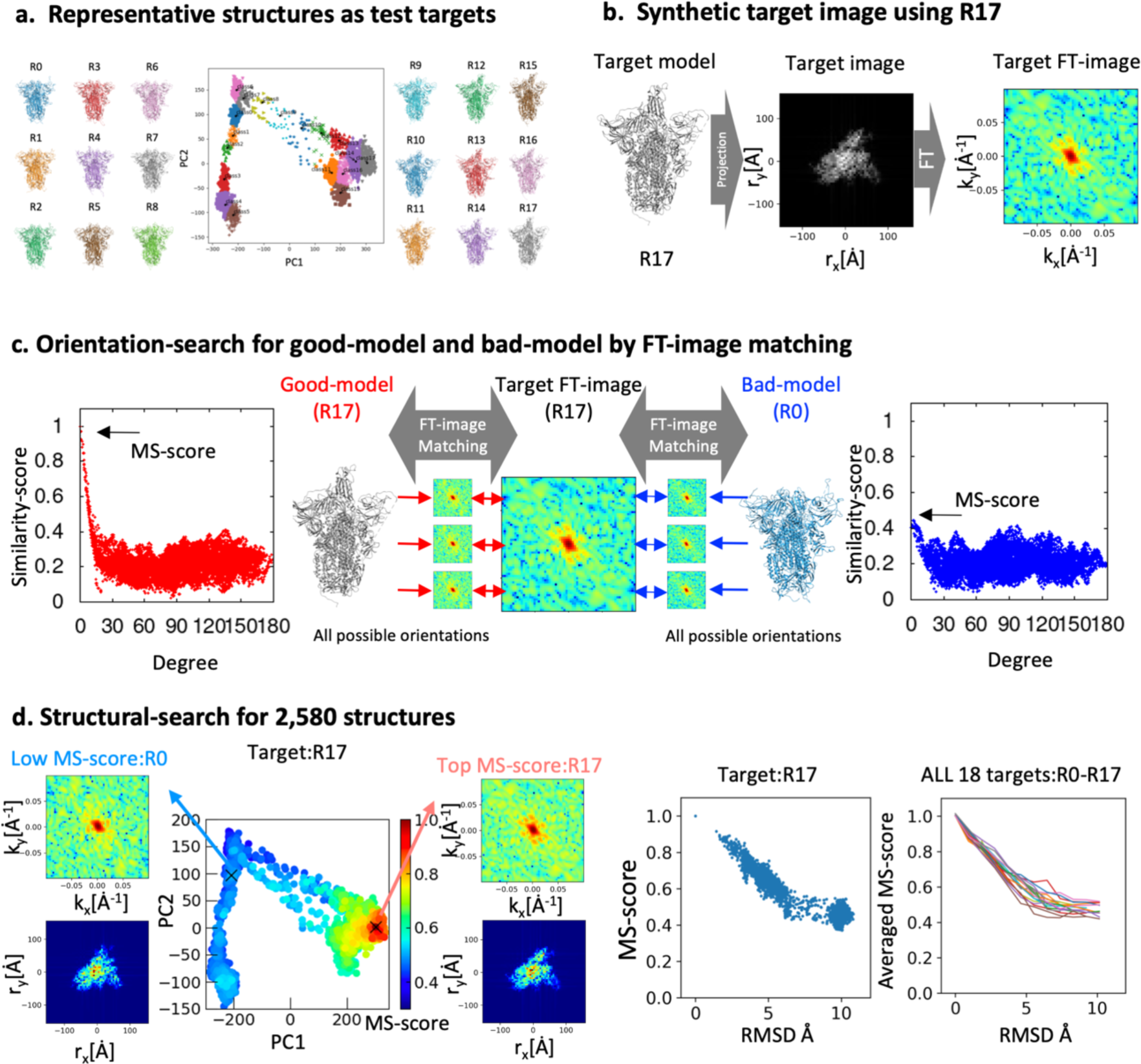
Test results of FT-image matching using synthetic data. (a) Clustering was performed in the structural space, selecting 18 representative structures from R0 to R17 as test target structures. (b) Using R17 as an example, a projection image was created along with the simulated target image and target FT-image. (c) An example of orientation search using two candidate structures: Good-model (R17) and Bad-model (R0) for all possible orientations. (d) An example of structural search for all candidate structures (2,580 structures). It can be confirmed that high MS-scores are present near Good-model (R17). A similar trend was observed when examining the dependence on RMSD from the target structure.

We created 2D projection images from these 18 representative 3D models at random orientations, which served as the target test single-particle images (Figure 2b). By conducting FT-image matching between the synthesized all-around single-particle images created from the same structure (good-model, R17 in the PCA plot) and the target test images, we obtained a similarity score distribution that exhibited a prominent peak value (Maximum similarity score, MS-score) corresponding to a specific orientation (Figure 2c). The peak position matched the orientation of the target test image (Figure 2d). This confirms that the FT-image similarity score can be used as an indicator to estimate the correct orientation.

Next, we performed orientation searching using a structure that differs from the target structure (bad-model, R0 in the PCA plot). In contrast to the results with the good-model, no significant peaks were observed, and the best score was below 0.5, significantly lower than the MS-score of 1 obtained with the good-model (Figure 2d). Furthermore, when examining the relationship between the RMSD value, which represents the deviation from the target structure, and the MS-score, we found that as the RMSD value increased, the MS-score decreased (Figure 2d). This indicates that using the MS-score allows for the estimation of the correct structure.

The successful estimation of the correct 3D structure at the correct orientation based on these similarity scores was achieved for all 18 target test images (See SI1). This demonstrates the feasibility of conducting single-image analysis to estimate a plausible 3D structure from a 2D single-particle image using the MS-score obtained from FT-image matching.

### Experimental CryoEM Single-Image Analysis of the SARS-CoV-2 Spike Protein

Validation using simulation data demonstrated that single-image analysis can be conducted through FT-image matching, leading to the application of the cryoMDM method on experimental cryoEM images. We performed cryoEM measurements targeting the spike protein, picking up 19,3125 single-particle images from cryoEM micrographs [31] and compressing the information into a 12-dimensional latent space to train the VAE model (See method).

As a result, the single-particle images generated by the VAE decoder showed a significant improvement in quality, with an average S/N ratio approximately 1.8 times higher while retaining the features of the original single-particle images (Figure SI2). We then conducted global and local searches in the structural space using these noise-improved single-particle images. In this analysis, we used representative images from 26 clusters that are narrowly distributed out of the 38 clusters obtained through image clustering in the image space as the target images for cryoMDM (See SI3).

Observing the distribution of MS-scores obtained from the global search for a representative target image (T4) revealed the presence of structural regions that yield significantly high MS-scores (Figure 3a: Global search). Among these, the structure with the highest MS-score (Promising structure) featured one RBD in an uplifted state, and the similarity score distribution with respect to molecular orientation showed a prominent peak at around 30 degrees (Figure 3a: Orientation search). Additionally, the wave number space image and real space image at this peak position (Figure 3a: Top MS-score images) were both found to closely match the experimental image. “We successfully confirmed a similar trend for all experimental images with 26 cases in the global search (see SI4).

**Figure 3.**
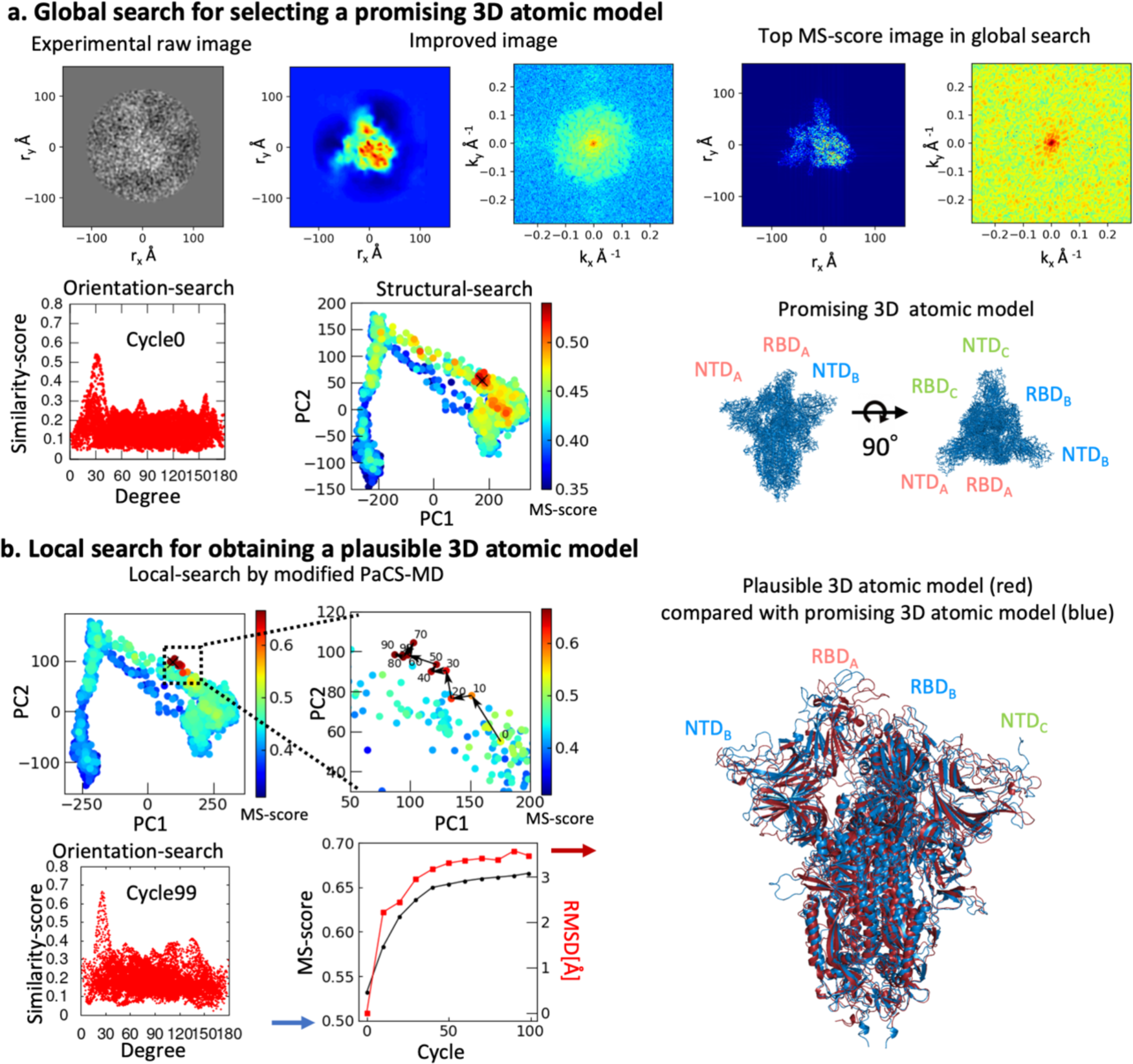
Example of cryoMDM analysis using cryoEM experimental single-particle image. (a) Example of global structural search targeting the cryoEM experimental image (T4) of the spike protein. Compared to the raw cryoEM experimental images, the improved images have enhanced image quality. Results of structural search are shown, with the position of the highest MS-score structure indicated by an “X.” The top MS-score image and 3D atomic model in the global search are also displayed. (b) Example of local structural search. Left panel: The transition diagram of MS-scores during local search (from cycle 0 to cycle 100) is presented. Additionally, the cycle dependence of MS-scores and RMSD is also shown. Results of orientation search for cycle 0 and cycle 100 are also shown. Right panel: Comparison of plausible structure and promising structure estimated from the T4 target image. In this example, the domain motions of RBD and NTD were observed in the local search.

Next, we performed a local structural search using the Modified PaCS-MD method, with the Promising structure from the global search as the initial structure and the MS-score as a constraint (Figure 3b). Looking at the MS-score map and its enlarged view, we observed that the search was exploring neighboring structures while generating a structure different from the initially prepared one starting from the Promising structure. Furthermore, examining the trend of the MS-score over the structure generation cycles revealed that the MS-score increased from about 0.53 in cycle 0 to approximately 0.66, indicating that a structure more consistent with the target image was being generated, as shown in the Top MS-score image.

When confirming the structure yielding the Top MS-score image in the local search, we found that the RBD_A_ in the Open state observed in the Promising structure was transitioning closer to the RBD_B_ state.

### Elucidation of Intermediate states of the SARS-CoV-2 Spike Protein

By comprehensively conducting both global and local structural searches on the experimental single particle images present in the electron microscope micrographs, we anticipate elucidating the diverse structural states that are averaged out and lost in experimental conditions.

Accordingly, we similarly performed global and local structural searches using FT-image matching on the 26 representative images, obtaining the plausible 3D structures corresponding to each individual single particle image (Figure 4). Since all these structures are derived as 3D atomic models, detailed analyses of each structural state can be carried out.

**Figure 4.**
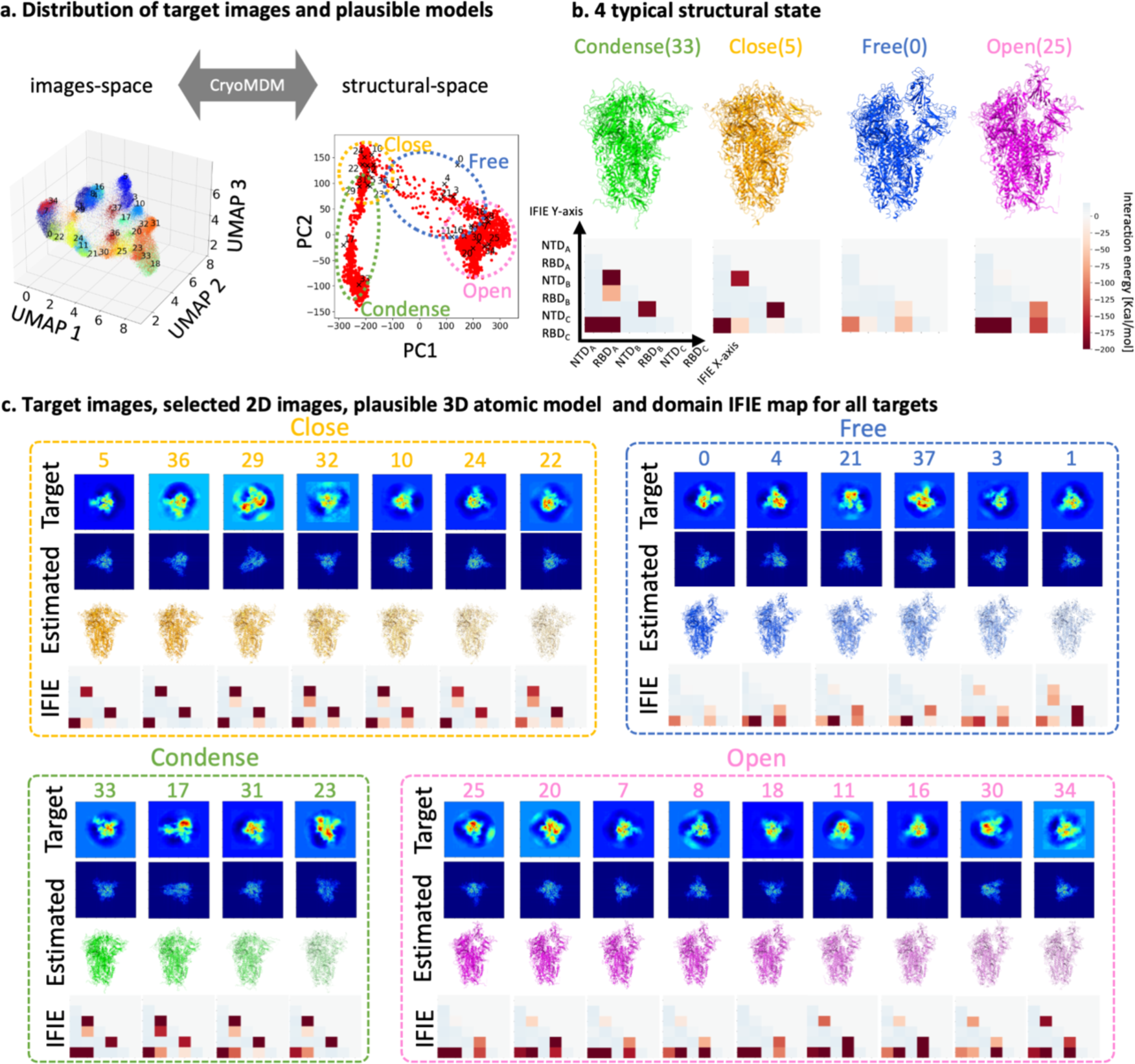
(a) Left: Distribution of images in the VAE latent space (image space). Only points belonging to the target image cluster are shown. Right: Results of the two-stage structural search (global-search and local-search) for all target images (a total of 26 target images). The plausible structures in the structural space are indicated by ‘×’ marks. (b) Structures of the 4 typical structural states (Condense, Close, Free, Open) along with their Domain IFIE maps. (c) Domain IFIE maps for each target image and the estimated structures (selected 2D images, plausible 3 D atomic models).

In this study, we performed quantum chemical calculations using the Fragment Molecular Orbital (FMO) method[32] on the obtained 3D atomic models, conducting domain IFIE analysis, which allowed us to classify the structures into four characteristic states (Condense, Close, Free, Open) by IFIE patterns, including previously unobserved metastable structural states (Figure 4, SI5). The typical domain IFIE patterns are shown in the figure4b for each states. Furthermore, we identified the molecular interactions contributing to the stabilization of each state at the residue level (Figures SI6, SI7, SI9, and supplementary text).

Interestingly, the obtained structures included not only the Close state (T5, 22, 29, 32, 24, 10, 36) and Open state (T25, 20, 7, 8, 30, 34, 11, 18, 16) that have already been reported through experimental approaches, but also two new structural states that had not been captured experimentally: the Condense state, where the RBD is in close proximity to surrounding domains (T33, 17, 31, 23), and the Free state of the RBD (T0, 4, 3, 21, 1, 37). These newly discovered structural states are thought to represent intermediate states that appear between the continuous structural changes of the Open and Close states in a non-bound state.

### Evaluation of Diverse Structural States in Multi-Image Analysis

While novel multiple conformational states were revealed by the cryoMDM method, it is crucial to verify whether density maps consistent with these structures can be reconstructed from actual experimental images. Therefore, based on the image group with MS-scores exceeding a certain threshold obtained through cryoMDM, we reconstructed 3D density maps using standard procedures and compared them with the 3D structures derived from cryoMDM to validate the robustness of this method.

When reconstructing a 3D density map with a resolution above a certain level using standard procedures, it typically requires tens of thousands to hundreds of thousands of single-particle images. However, as the number of images increases, flexible regions can lose their unique characteristics due to averaging. Therefore, to conduct this validation, we examined how to determine an image set size that effectively captures the uniqueness of the density map while still allowing for observable characteristics.

Using 128,739 single-particle images obtained through standard structural determination procedures, we reconstructed a 3D density map with an overall resolution of 3.8 Å, comparable to previously reported quality. By applying a threshold to the MS-score and selecting the top images, we created down-sampled image sets of 77,734 (threshold: 0.45), 39,922 (threshold: 0.5), 15,131 (threshold: 0.55), and 7,943 (threshold: 0.57) images, and conducted 3D reconstructions for each. The overall resolutions obtained were 4.0 Å, 4.7 Å, 6.8 Å, and 8.3 Å, respectively (Figure. 5a). This indicates that the final overall resolution of the spike protein depends on the number of images belonging to similar classes.

**Figure 5.**
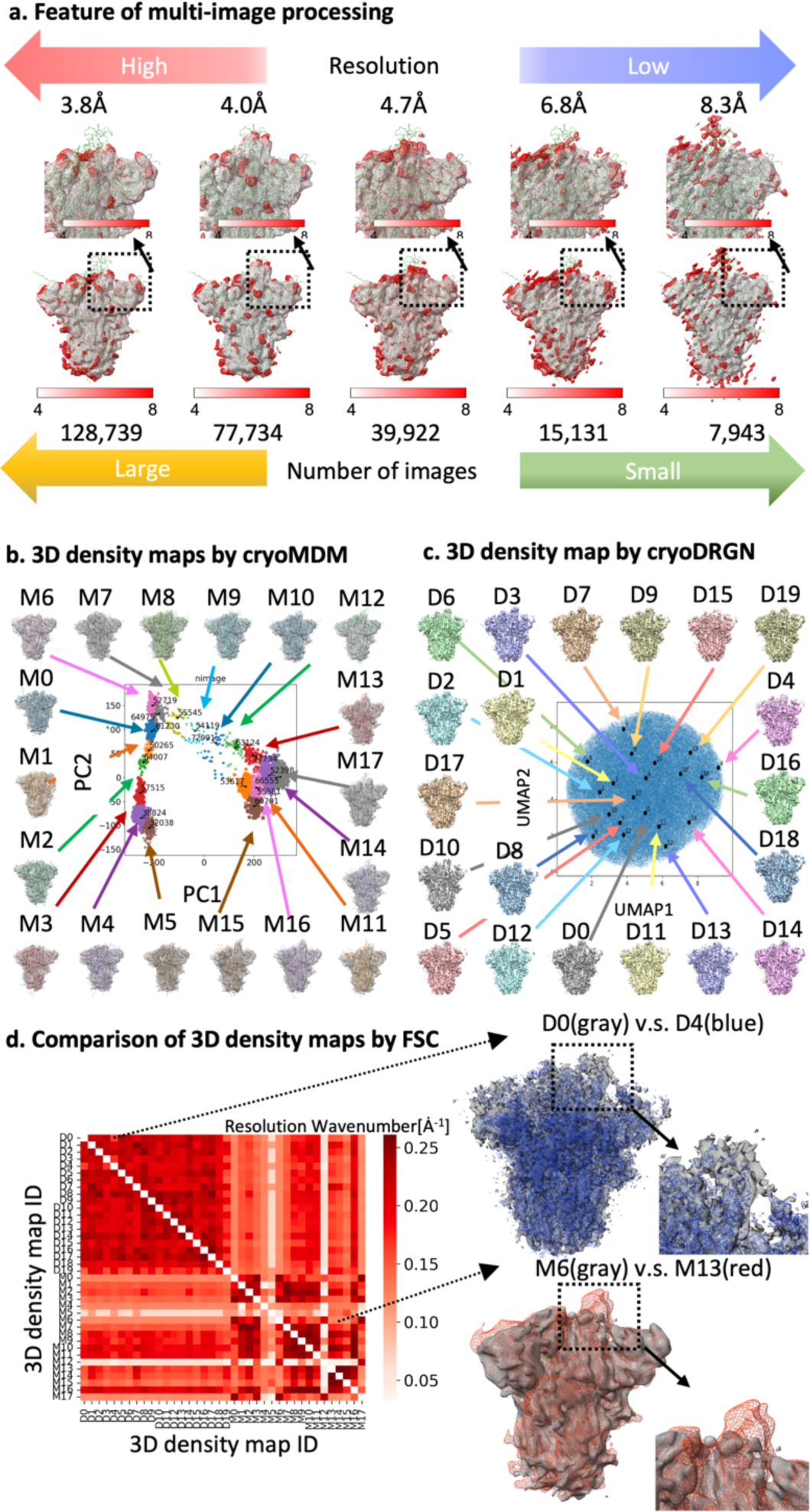
(a) The dependence of the number of similar images on the 3D reconstruction obtained through Multi-Image Analysis. (b) The 3D density maps constructed using cryoMDM (M0-M17). (c) The 3D density maps constructed using cryoDRGN (D0-D19). (d) Comparison of the 3D density map groups using FSC. The resolution wave number at which FSC reaches 0.5 is displayed as a heatmap. Additionally, the structures where the differences in the density maps are maximized are displayed (M6-M13 and D0-D4).

At the maximum of 128,739 images, it was observed that the information from flexible structural regions, particularly around the RBD, became less visible. Therefore, we decided to adopt an image set that achieved a resolution of 4.0 Å (threshold: 0.45), which was more effective in expressing the uniqueness of the density map while still allowing for observation. We conducted FT-image matching using the diverse 18 representative structures and 193,125 VAE target images (193,125 target images × 18 representative structures × 10,000 orientations = 36 billion comparisons). For each representative structure, we selected VAE target images showing a similarity score of 0.45 or higher, resulting in approximately 60,000 images for each representative structure. Using these VAE target images, we performed multi-image analysis to achieve 3D reconstructions, successfully obtaining 3D density maps with a resolution of about 4 Å (Multi-image analysis M0-M17) (Figure. 5b).

When comparing the resulting 3D density maps with the atomic models used for image selection, it was clear that, despite some ambiguity originating from multi-image analysis, the 3D density maps were consistent with the 18 diverse structural states obtained by the cryoMDM method. For example, the atomic models used for image selection for M6 and M15 showed the RBD region in the Open and Close states, respectively. Similarly, the RBD region in the reconstructed 3D density maps based on these atomic models also displayed Open and Close states. This result indicates that the FT-image matching through VAE images is functioning effectively, strongly supporting the validity of the cryoMDM approach.

### Comparison of cryoMDM with Multi-Image Analysis methods

CryoDRGN [16, 33, 34] is widely used as a multi-image analysis method to obtain multiplu conformational states from single-particle image data set. To compare the ability to capture polymorphisms between cryoMDM and cryoDRGN, we conducted an analysis using the spike protein dataset (19,3125 images) employed in this study (See SI10). Based on the frequency of empirical distributions in the latent space, we obtained 20 representative 3D density maps (D0-D19) (Figure 5c). All the density maps obtained from cryoDRGN exhibited structures with one RBD raised, indicating that cryoMDM is more sensitive in capturing minor structural states. We quantitatively evaluated the similarity between the 3D density maps obtained from cryoMDM and cryoDRGN using Fourier Shell Correlation (FSC). The results showed that the density maps derived from cryoDRGN (D0-D19) were similar to each other, while the density maps obtained from cryoMDM (M0-M17) exhibited high structural variation (Figure 5d). Furthermore, comparing D0-D19 with M0-M17 revealed that certain maps, such as M8-M11 and M16, closely resemble those obtained by cryoDRGN. This indicates that cryoMDM captures not only the conformational states identified by existing AI methods but also entirely different conformational states.

### Construction of cryoEM image trajectories

In MD simulations, it is possible to capture structural changes of proteins over time. The cryoMDM method can associate each experimental image with a 3D structure that contains time information, allowing us to add a time axis to cryoEM single-particle images (Figure 6a). From the clustering analysis conducted in the VAE latent space, we selected images belonging to cluster 33 as target images and added time information to each single-particle image through FT-image matching with the global structural space. By rearranging the single-particle image set according to the added time information, we created a video of the cryoEM experimental single-particle images, which shows that the three RBDs transition through various structural states over time (See Figure 6b, Video SI11). This result suggests that the time information lost during sample freezing can be recovered through cryoMDM, and the observed dynamic behavior reflects the inherent ‘flexibility’ of the spike protein in the real world.

**Figure 6.**
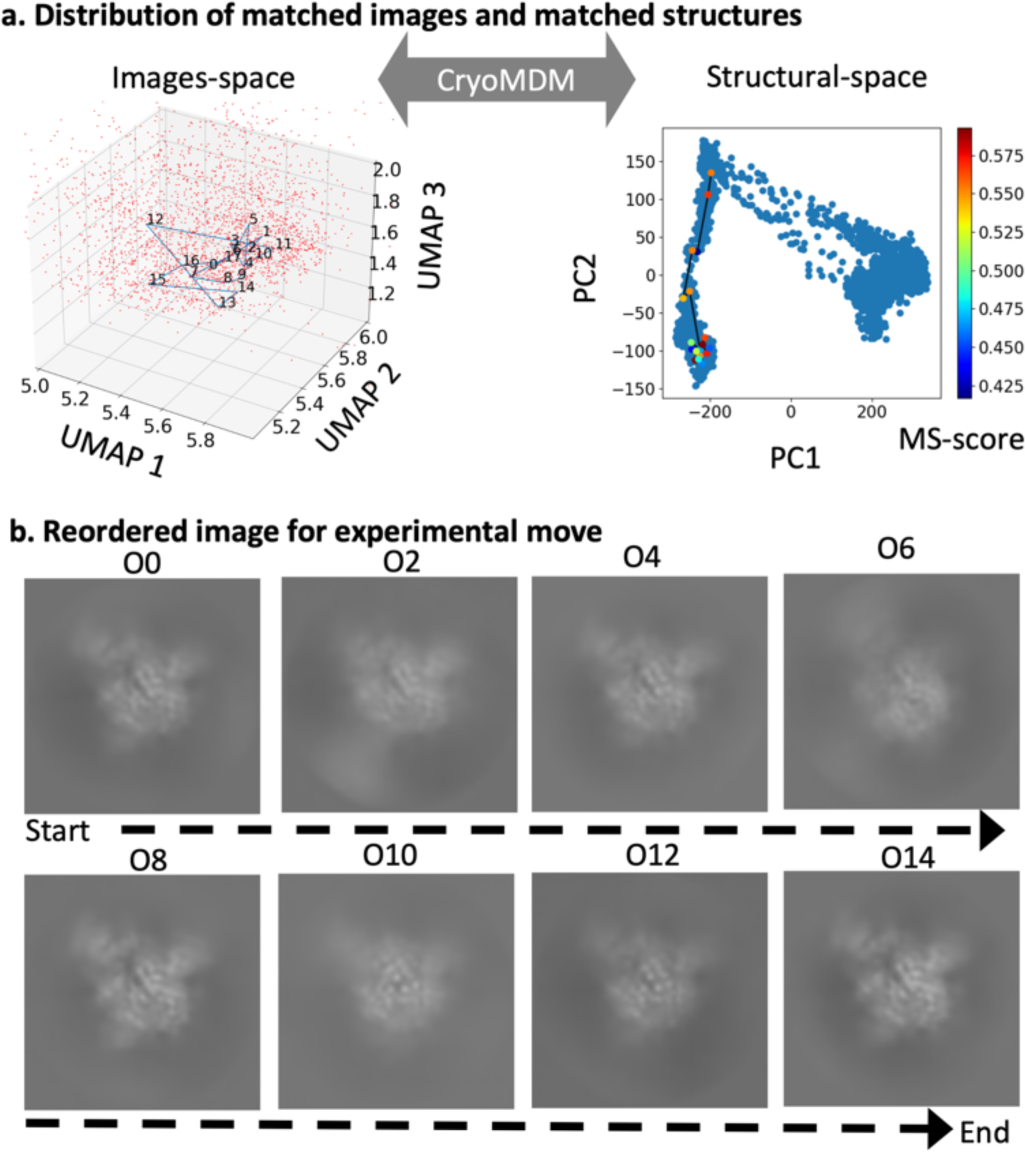
Procedure for constructing cryoEM image trajectories. (a) Left panel: Trajectories of images in the image space linked by cryoMDM. Right panel: Trajectories of structures in the structural space linked by cryoMDM. (b) Experimental images reordered along the MD time axis. https://riken-share.box.com/s/t4yskzq6hwxpgjcvvijhy075ygzzkq3f

## Discussion

Research has progressed in capturing the continuous structural changes of biomolecules by utilizing experimental measurements and molecular simulations. Experimental measurements allow for the assessment of various conformational states of biomolecules by adjusting the sample. However, the physical quantities obtained are often lower-dimensional than the target being measured, and low S/N ratios can pose problems. While statistical analysis can complement these characteristics, averaging and other processes can lead to a loss of the individuality reflected in the individual measurement data. On the other hand, molecular simulations of biomolecules can reproduce the time-dependent changes in 3D structures with high resolution down to the microsecond scale. However, large structural changes that occur between metastable states are probabilistic and happen as rare events on the millisecond scale. As a result, standard MD simulations often fail to capture all diverse structural changes, leading to the proposal of various methods, such as multi-canonical methods[35] and replica exchange methods[36], to efficiently sample the structural space of proteins. To elucidate the diverse structural states of biomolecules, it is essential to integratively utilize experimental measurements that, while lower in resolution, include various structural states, alongside simulations that can yield high-resolution structures. From this perspective, this paper proposes the cryoMDM method, a simulation-driven structural estimation technique that comprehensively estimates various atomic 3D structural models from noisy cryo-electron microscopy single-particle images through the integration of experiments, AI, and simulations. By constructing a structural estimation workflow consisting of four procedures, including image quality enhancement through VAE, on a supercomputer, we have realized single-image analysis that estimates a single 3D atomic model from a single particle image.

The cryoMDM method is a highly accurate structural estimation technique that respects the individuality of experimentally observed data and is applicable to all biomacromolecules. This method has three key advantages:

Advantage 1: It is possible to directly estimate 3D atomic models to evaluate various intermediate states from a single-particle noisy image that exhibits individuality.

Advantage 2: By using the estimated all-atom models, detailed structural analyses can be performed, leading to a better understanding of molecular mechanisms.

Advantage 3: The cryoMDM method allows for the addition of temporal information, which is typically lost in cryoEM experiments.

In this study, we identified four characteristic structural states of the spike protein based on advantages 1 and 2, and elucidated the residues contributing to their stabilization through quantum chemical calculations. These structural states include metastable states that are difficult to capture experimentally, which could deepen our structural understanding of the molecular mechanisms involved in the entry of the novel coronavirus into cells.

Previous research has demonstrated that minor structural states are crucial for the functional regulation of biomacromolecules. Additionally, intrinsically disordered regions lacking specific structural states are particularly important for the functional expression of biomacromolecules in eukaryotes. While capturing these structural states experimentally is generally challenging, the single-molecule analysis method cryoMDM has the potential to significantly advance our structural understanding of these states.

Furthermore, through advantage 3, we can directly observe the dynamic movements of regions such as the RBD and tail, rooted in cryoEM experimental data. Until now, capturing the structural states of biomacromolecules with such high spatiotemporal resolution required advanced instruments like high-speed atomic force microscopy or XFEL. The cryoMDM method paves the way for utilizing cryoEM not only as a powerful structural determination technique but also as a means of analyzing biomacromolecules with high temporal resolution. In this study, we demonstrated that the cryoMDM method captures a greater diversity of structural states compared to the widely used cryoDRGN method (Figure 5d), indicating that cryoMDM allows for more effective observation of structural polymorphism than traditional methods. While cryoDRGN uses three-dimensional orientation information from each 2D image as prior information to express polymorphs as 3D density maps, this results in 3D density maps that inherit the characteristics of multi-image analysis. Consequently, the issue of averaging leads to a loss of clarity in flexible regions.

In contrast, the cryoMDM method derives an atomic model from a single particle image rather than an averaged 3D density map, allowing for a more nuanced representation of diverse structural states.

On the other hand, the cryoMDM method currently faces three limitations.

Limitation 1: Uncertainty in estimation accuracy dependent on image noise. Limitation 2: Trapping in local minima during structural search.

Limitation 3: High computational cost associated with extensive structural sampling through large-scale matching and molecular dynamics calculations.

To address Limitations 1 and 2, this study employs variance as an indicator to monitor the convergence of the Modified PaCS-MD. By ensuring that the variance stabilizes to a certain level, we can determine when to terminate local search. Regarding Limitation 3, the computational cost for a single target image (1 target image × 2,580 structures × approximately 10,000 orientations) in this study is approximately 215 NH (using 323 nodes). If the number of target images is 10,000, the required computational cost would amount to around 40,850,000 NH.

To mitigate this impractical computational cost, this study successfully reduced it to 5,590 NH by selecting 26 representative target images based on image clustering in the VAE latent space. The fact that the 3D density maps reconstructed using the selected images align well with the atomic models used for image selection strongly supports the validity of this approach. As cryoEM technology continues to advance and HPCI environments are further developed, we anticipate that these limitations will be resolved, leading to improved spatiotemporal resolution and accuracy of the cryoMDM method.

In this study, we developed a novel data assimilation method called cryoMDM, which estimates multiple conformational states from single-particle images obtained through cryoEM measurements in a simulation-driven manner. The cryoMDM method requires only one reliable biomolecular 3D structure model and cryoEM images as input, making it highly versatile. By combining the estimated structures from AlphaFold2[37] with MD simulation, we expect to establish a pipeline that elucidates multiple conformational states based on sequence information and cryoEM images.

Furthermore, by applying this method to cryoEM tomography data, it will be possible to visualize diverse instantaneous structures within a cellular environment at high resolution, with one image representing one structure. This capability is anticipated to bridge the gap between cell biology and molecular biology, contributing to innovations in life sciences and drug discovery processes.

## Method

### cryoMDM method

In cryoEM single-particle structure analysis, electron beam is directed at a frozen sample, and 2D images are captured using a 2D detector. The field of view contains multiple molecules, resulting in images of molecules at various orientations. To minimize sample damage, the signal-to-noise (S/N) ratio of the obtained 2D images is typically low. Generally, numerous 2D images from various molecular orientations are combined to construct an average 3D structure. However, biomolecules are flexible and can adopt multiple conformational states.

On the other hand, the cryoMDM method supplements observational data with simulation-derived data, allowing for the high-resolution estimation of biomolecular structures from limited observational data. This method employs “structural sampling” and “FT-image matching” for structural search. To improve efficiency, structural search is conducted through global and local searches.

In the global search phase, many candidate structures are hypothesized, and for each candidate structure, a multitude of candidate images considering the degrees of freedom in molecular orientation are prepared. The candidate structures can be derived from various structural databases, ab initio modeling, and molecular dynamics (MD) simulations. Similar candidate images to the target image are explored using FT-image matching, and promising structures are selected.

Next, local searches are conducted to refine the structure around the promising candidates. If a candidate image closely resembles the target image, the candidate image can be linked to the candidate structure, allowing for the estimation of the 3D structure. This enables the extraction of 3D structural information from a single target image. By performing similar analyses on individual target images that exhibit structural uniqueness, the method has the potential to elucidate multiple conformational states.

### FT-image Matching

CryoEM images are obtained by using an electron beam as the probe light to create real-space images. To search for similar patterns in real space, it is necessary to consider translational and rotational degrees of freedom. Therefore, in this study, search is conducted in wavevector space. By performing a Fourier transform on the real-space images and calculating their intensity, the wavevector space representation of the target image is prepared. Additionally, various wavevector space images (candidate images) at different orientations are generated from the candidate structures. The correlation value c_ij_(ξ, α) between a pair of wavevector space images I and j is defined as follows.

This method allows for the normalization of intensity images based on the wavevector dependence of the expected value, which helps capture correlations over a broader wavevector range, enabling the effective utilization of even weaker signals as valid data.

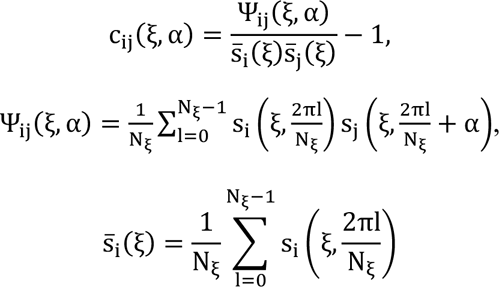

The rotation of the molecule with respect to the incident beam axis appears as a rotation around the center in wavevector space images. By considering this rotational degree of freedom α, the above correlation coefficient allows us to simultaneously obtain correlation values for patterns rotated 360 degrees in the two-dimensional plane. The correlation values expressed as functions ofα and 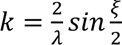 are referred to as the correlation map. Here, *ξ* = 2*θ* represents the scattering angle. Additionally s denotes the wavevector space intensity of the observed image, *s* represents the expected value of the intensity, λ is the wavelength of the incident probe light, and *N*_*ξ*-_ indicates the discrete number of pixels along the concentric circles.

Furthermore, we define the integrated correlation value (or integrated correlation map) I_c,ij_(k_up_, α) by integrating the correlation value c_ij_(ξ, α) over the directions of α and k. This integration helps reduce the influence of noise.

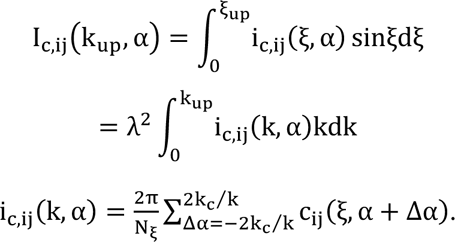

λ represents the wavelength of the incidentbeam. k_c_ is the correlation length of the speckle pattern, which is on the order of the molecular size. Additionally, k_up_ denotes the maximum wavevector of the wavevector space image used for similarity determination calculations. In the integrated correlation map, we define the direction of the correlation line as 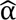, and the maximum integrated correlation value when viewed in the k direction at 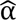 is denoted as 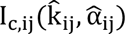. By normalizing this with the self-correlation terms of each wavevector space image obtained in a similar manner, we quantify the similarity of the patterns of a pair of wavevector space images as a normalized matching score 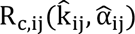: (referred to as the similarity score).

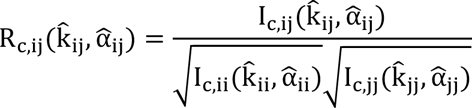

Through the wavevector space images, we estimate the similarity between the target structure and candidate structures. However, even for the same candidate structure, the patterns of the wavevector space images differ depending on the orientation of the incident probe light (molecular orientation). Therefore, to explore similar structures using “wavevector space matching,” it is necessary to simultaneously identify similar molecular orientations for the target image (molecular orientation matching). For each candidate structure, we prepared wavevector space images for 10,240 different orientations. The highest R_c_ value obtained from molecular orientation matching is defined as the structural agreement (Maximum Similarity Score; MS-score) for that candidate structure.

In the experimental data analysis of the spike protein, FT-image matching was performed using VAE images of size 128x128 pixels (with a pixel size of 1.76 Å) following these steps. To mitigate the effects of aliasing during the Fourier transform, the pixel average was padded, converting the image to 180x180 pixels and creating wavevector space images. Subsequently, the similarity scores were calculated for the wavevector space images focused on the central region of 64x64 pixels, where the signal patterns were present.

### Preparation of the Global Structural Space Using Public Data and Selection of Promising Structures via Global Search Workflow

To select promising structures for each target image using FT-image matching, a global search workflow was established on “Fugaku.”

In this global search, a pool of candidate structures was prepared in advance. It is desirable for the candidate structures to cover a wide structural space while maintaining protein-like conformations. Various methods can be considered for preparing candidate structures. In this study, we adopted a diverse set of candidate structures by combining publicly available all-atom MD data of the spike protein.

To prepare a diverse set of spike protein structures in the simulation space, we utilized the results of multiple MD simulations. In addition to all-atom MD simulations starting from the inactive Close structure (PDB: 6VXX)[38] and the active Open structure (PDB: 6VSB)[39] of fully glycosylated SARS-CoV-2 S-protein in solution, we integrated the results of Targeted Molecular Dynamics (TMD) simulations that connect the two structural states (Close/Open). A total of 2,580 snapshots were extracted every 1 ns from the publicly available trajectories listed below (1 to 10), resulting in a global structural space composed of 2,580 structures.

Normal AA-MD trajectory group:

1. Extracted 1,000 structures from MD1_Up (1ms MD from up)
2. Extracted 200 structures from MD2_Up (200ns MD from up)
3. Extracted 1,000 structures from MD1_Down (1ms MD from down)
4. Extracted 200 structures from MD2_Down (200ns MD from down) TMD trajectory group:
5. Extracted 20 structures from TMD1_toUp (20ns TMD from down to up)
6. Extracted 20 structures from TMD2_toUp (20ns TMD from down to up)
7. Extracted 50 structures from TMD3_toUp trajectory (50ns TMD from down to up)
8. Extracted 20 structures from TMD1_toDown trajectory (20ns TMD from up to down)
9. Extracted 20 structures from TMD2_toDown trajectory (20ns TMD from up to down)
10. Extracted 50 structures from TMD3_toDown trajectory (50ns TMD from up to down)

By conducting image matching between the target images and the candidate structure pool, promising structures were selected. In the FT-image matching, a complete set of approximately 10,000 2D projection images was prepared for each candidate structure, and their similarities were assessed to obtain the similarity value for the most promising molecular orientation as the structural consistency score. The structural consistency scores were calculated for all candidate structures, and the structure with the highest score was selected as the promising structure. As a result, the number of FT-image matching calculations in the global search was 38 target images × 2,580 structures × 10,000 orientations = 830 million calculations.

Regarding the computational cost of the global search, the cost for one target image (1 target image × 2,580 structures × approximately 10,000 orientations) was about 215 NH (using 323 nodes). Based on this, estimating the cost for 2,580 structures × 10,000 target images results in approximately 40,850,000 NH. Given the overwhelming computational demand for executing all targets, image clustering in the VAE latent space was performed to efficiently select target images for the analysis.

### Structure Refinement via Local Search Workflow

To efficiently explore structures around promising candidates for structure refinement, we developed a local search workflow. The underlying algorithm, PaCS-MD [40], is one method for efficiently conducting structure sampling that approaches the target, consisting mainly of two steps: 1) performing temperature-controlled MD in parallel using multiple selected structures, and 2) selecting higher structures based on an index that represents the deviation from the target. By repeating steps 1 and 2, we aim to explore structures that move closer to the target.

In this study, we adopted the similarity value between the experimental image and the candidate diffraction image set as the index representing the deviation from the target. While a candidate structure needs to be prepared in advance for global search, local search features the generation of candidate structure groups on the fly through MD in conjunction with image matching in a series of workflows. The MD engine can utilize various software, such as GROMACS[41], GENEISI[42], and CafeMol[43]. Here, we opted for all-atom MD (GROMACS), which allows for stable and direct estimation of the full atomic structure model.

Regarding the computational cost of local search, conducting PaCS-MD simulations with 120 parallel MDs (120 × 10 structures = 1,200 structures) over 100 cycles (using 151 nodes) consumes approximately 14,500 NH. In this case, 120,000 structures (120 parallel × 10 structures × 100 cycles) will be matched. The length of each MD cycle is 20 ps (10,000 steps, 1 step = 2 fs), resulting in a total of 2 ns of MD over 100 cycles. The resource consumption for conducting PaCS calculations with 120 parallel MDs (120 × 10 structures = 1,200 structures) over 100 cycles for each of the 26 target images is 14,500 × 26 = 377,000 NH.

### Protocol for the MD simulation in the PaCS-MD

To relax the initial system for PaCS-MD, energy minimization and MD simulations were conducted using modules of GROMACS v. 2019.1[44]by adopting the AMBER ff99SB force field[45], according to the following protocol. A target molecule was placed in a cubic space 10 Å larger than the target molecule in the x, y, and z directions and was solvated with 160 mM NaCl solution using the genion module of GROMACS. Energy minimization of the entire system was conducted using the steepest descent algorithm. Initial velocities were randomly generated according to the Maxwell-Boltzmann distribution at 300K. The production run of conformational sampling with a 20 ps MD simulation was conducted. The trajectory from 2ps to 20 ps every 2ps .i.e. 10 structures of the production run was used in subsequent analysis of the matching in PaCS-MD.

### Preparation of Image Data and 3D Reconstruction Using RELION

The cryoEM movies of the SARS-CoV-2 spike protein were processed using the previously reported data [31] and all image analysis was performed using RELION software [8]. After performing motion correction of movies and estimating CTF parameters, some particles were manually picked. The template images were generated through 2D classification for manual picked particles, and particles were subsequently automatically picked using template-based particle picking algorithm. The picked particles were extracted into a box of 360 x 360 pixels.

All images were classified into 5 classes by 3D classification, and a totally of about 190,000 particles in the 3 classes with clear structural map and distinct conformations were selected. By performing 3D auto-refine and post analysis for the images, a final structure with a resolution of 4 Å was reconstructed.

To cryoMDM analysis, all images were re-extracted from 360 x 360 pixels to 320 x 320 pixels and resized to 256 x 256 pixels and 128x128 pixels by Fourier shrink. Individual particle images were extracted as intermediate image files from the 2D classification process using a modified version of RELION software. VAE analysis was carried out using all individual particle images to improve the image quality.

### VAE Network Model and Learning Procedure

The VAE (Variational Autoencoder) encoder used in this study has four layers of 2D convolutional layers, as shown in FigureSI12, with the image size reduced by half at each layer. The shape of the 2D convolutional kernels is 3x3, and the stride is 2 in both vertical and horizontal directions. The number of kernel filters from the 1st to the 4th layer is 128, 256, 512, and 1024, respectively. Each 2D convolutional layer includes a dropout layer with a dropout rate of 0.3. The activation function used for the 2D convolutional layers is ReLU. The input image of size 128x128 is compressed to 12 dimensions by the encoder, and then returned to the size of 128x128 by the decoder, which has a structure nearly opposite to that of the encoder, as shown in Figure SI12. The variance of the random numbers used in the reparameterization trick [23] is set to 0.3.

The noise reduction and classification of the target images using VAE were conducted in the following steps. First, the target images were used as training data for VAE learning. The loss function employed was binary cross-entropy, and mini-batch learning was performed with 32 images per batch for a maximum of 100 epochs. Early stopping was applied, terminating the training if the minimum loss function value did not change for more than five consecutive epochs. Next, the trained model was used to transform the target images, resulting in VAE-generated images with reduced noise. Clustering of the VAE-generated images was performed using Gaussian Mixture Model (GMM). The clustering was conducted with multiple class numbers (ranging from 1 to 50), adopting the class number that yielded the highest Bayesian Information Criterion (BIC). Dimensionality reduction and visualization were performed using UMAP (Uniform Manifold Approximation and Projection for Dimension Reduction, [46, 47]. The 12-dimensional latent space data obtained from the VAE encoder for the target images was compressed to two dimensions using UMAP and visualized based on the class labels obtained in step 2.

### Ensemble fragment molecular orbital calculation

To stably evaluate the estimated structures through the FMO method[48], ensemble FMO calculations were performed using the top 10 structures from the aforementioned local search. This approach helps reduce estimated noise and is expected to capture more intrinsic structural features. Specifically, from the results of the 100 cycles of local search, the top 10 structures with the highest scores were extracted from the cycles that included the structure with the highest score for each target. Since there are 26 types of targets, a total of 260 structures were obtained.

In conducting quantum chemical calculations for these 3D atomic model structures, a multi-step optimization was performed to optimize structural distortions and hydrogen positions.

Specifically, classical mechanics modeling using Amber99SB-ILDN[49] was employed for structural optimization calculations on the 260 structures with Gromacs version 2022.3. The Steepest Descent method was used as the calculation technique, applying a position constraint of 1000 kJ mol^(#^ nm^(%^ on the heavy atoms of the protein. The convergence conditions were set such that the maximum force was below 500 kJ mol^(#^ nm^(#^ for the first step and below 100 kJ mol^(#^ nm^(#^ for the second step. Through this process, optimized structures were obtained, free from local distortions.

To assess the validity of the optimized structures, the RMSD (Root Mean Square Deviation) of all atoms was calculated for each target. The structure with the highest score for each target served as the reference structure, and the RMSD was calculated for the remaining nine structures. The average and standard deviation of the RMSD within each target were plotted, as shown in SI7. As can be seen from the figure, the deviations between structures within the cluster were found to be within a maximum of 2.0 Å. Therefore, the top 10 structures within each target can be considered to possess structurally similar features.

For each structure obtained from local search and structural optimization, electronic state calculations were performed using the ABINIT-MP Open Ver. 1 Rev. 23Q2 software [50]with the Fragment Molecular Orbital (FMO) method. In the FMO approach, the system is divided into subunits called fragments, and the electronic energy is approximated using the values obtained from calculations for each fragment.

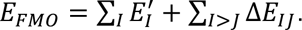

In the equation, E’I represents the energy of fragment I after subtracting the contribution from the environmental electrostatic potential, while DE_IJ_ is the interaction energy between fragments I and J obtained from the dimer energy. By expanding to trimers and tetramers, the FMO energy approaches the true energy of the system.

In this study, the FMO expansion was limited to dimers, and the electronic energy was calculated using the Hartree-Fock + MP2 method. The 6-31G(d) basis set was employed, and fragmentation was performed to ensure that, in principle, each fragment contained one residue (with the exception that pairs of cysteine residues cross-linked by disulfide bonds were treated as a single fragment). For dimers with inter-fragment distances equal to or greater than twice the sum of their van der Waals radii, the dimer-ES approximation was used. This represents the standard setting for contemporary FMO calculations. As a result, FMO calculations converged for a total of 245 structures. Following this, fragment interaction energy (IFIE) analyses were conducted on these results.

Specifically, the analysis of fragment interaction energies was conducted using PIEDA (pair-interaction energy decomposition analysis)[51]. In PIEDA, the energy decomposition method is applied to split the fragment interaction energy into four components:

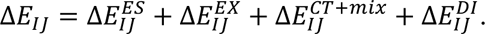

ES, EX, CT+mix, and DI represent the electrostatic interaction energy, exchange interaction energy, charge transfer energy, and dispersion energy, respectively, with all except ES being short-range interactions.

In this study, after averaging the energy components of fragment interactions obtained from the PIEDA analysis for each target, fragment pairs were defined as important fragments if the absolute value of any of the three components (short-range interactions) other than ES was 3 kcal/mol or greater.

Additionally, the plot showing the average and standard deviation of PIE (= ES + EX + CT + mix + DI) is presented in SI8.

## Data availability

The generated MD data are publicly available via the BSM-## (https://bsma.pdbj.org/entry/58). The cryoEM experimental movies are found in https://clinfo.med.kyoto-u.ac.jp/si/XX.XXXX/XXXXXX/.

## Acknowledgements

This work was supported by the FOCUS Establishing Supercomputing Center of Excellence project; MEXT as “Program for Promoting Researches on the Supercomputer Fugaku” (Simulation-and AI-driven next-generation medicine and drug discovery based on “Fugaku”, JPMXP1020230120).

This work used computational resources of the supercomputer Fugaku provided by RIKEN Center for Computational Science through the HPCI System Research Projects (Project IDs: hp220078, hp230102, hp230216, hp240109, hp240162, ra000018); the supercomputer system at the information initiative center, Hokkaido University, Sapporo, Japan through the HPCI System Research Projects(Project IDs: hp220078, hp230102) . Dr. Yasumasa Joti and Dr. Motohico Matsuda for their valuable input and insightful discussions during the development of this manuscript.

## Author contributions

A.T., and Y.O. designed the study. A.T. and Y.A implemented the cryoMDM workflow on Fugaku. T.K. conducted experimental measurements of cryoEM. T.K., S.M., A.T., and Y.S. conducted cryoEM data analysis. A.T., Y.S., and Y.A., performed the cryoMDM calculations on Fugaku. A.T. wrote the manuscript. All authors discussed the research, edited the manuscript, and approved its final version.

## Supplementary information

The Supplementary Information is available free of charge at https://XXX. Details of the supplementary results and methods, i.e., IS1.Results of structural search for 18 types of synthetic data, IS2.SNR analysis, IS3.Image distribution in a 12-dimensional latent space, IS4.Results of globule-search for all experimental target image, SI5. Four typical structural states estimated by the cryoMDM method, SI6: Results of FMO calculations for the four typical structural states, SI7. The RMSD plots of the top 10 scored structures, SI8. Results of PIEDA analysis, SI9: List of important interaction residues, SI10. Conducting of cryoDRGN analysis, SI11.Experimental movie, SI12. VAE Network Model.

