## Supplementary information for "CryoMDM; Molecular simulation Driven structural Matching Approach to Estimate Variety of Atomic Three-dimensional Model Based on Noisy cryoEM Single Particle Images"

**SI1:Results of structural search for 18 types of synthetic data**

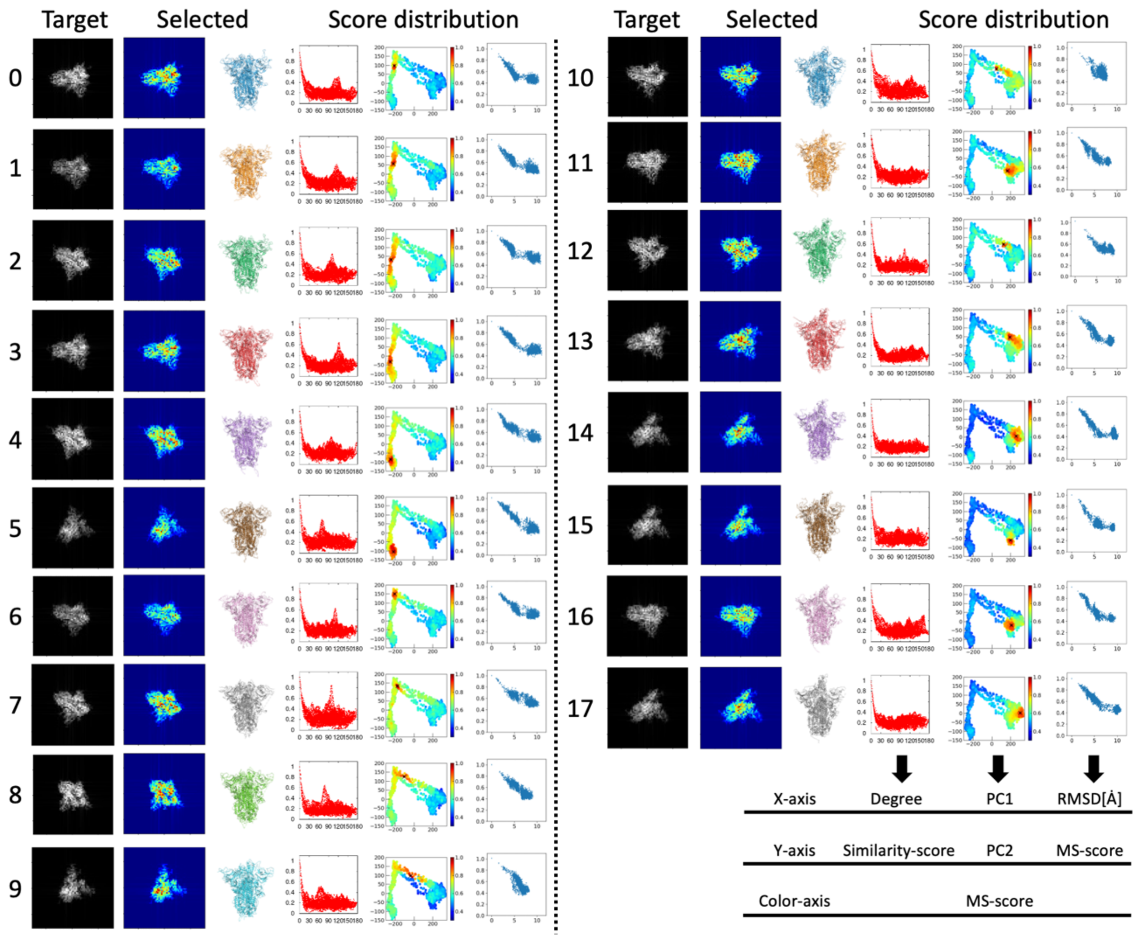


Figure SI1. Results of structural search for 18 types of synthetic data. Structural search was conducted for the 2,580 structures prepared for global structural search. Target: target images. Selected: selected 2D images and 3D structures from the search results. Score distribution: (left) Results of molecular orientation search when the MS-score is maximized. X-axis: Degree, Y-axis: Similarity score; (middle) MS-score distribution. X-axis: PC1, Y-axis: PC2; (right) RMSD dependence of MS-score. X-axis: RMSD, Y-axis: MS-score.

**SI2 : SNR analysis**

The signal-to-noise ratio (SNR) for the 19,3125 particle images used in VAE training was calculated for both the raw images and VAE images using the following equation

$$SNR[db] = 10log10(\mu^2/\sigma^2)$$

In the above, μ represents the mean pixel value and σ represents the standard deviation of the pixel values. Considering the image preanalysis done by RELION, μ and σ were calculated within a masked area on a circle.

The plot shows SNR (Raw) on the X-axis and SNR (VAE) on the Y-axis. In both cases, SNR (VAE) shows higher values, indicating that the image quality has improved due to VAE analysis. The maximum value of SNR(pred)/SNR(raw) was 2.53, the minimum was 1.23, and the average was 1.84.

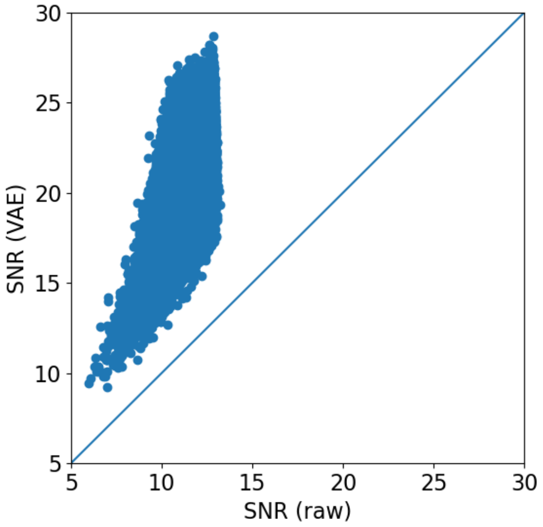


Figure SI2. SNR Distribution Plot. X-axis: SNR of raw images. Y-axis: SNR of VAE images

**SI3: Image distribution in a 12-dimensional latent space**

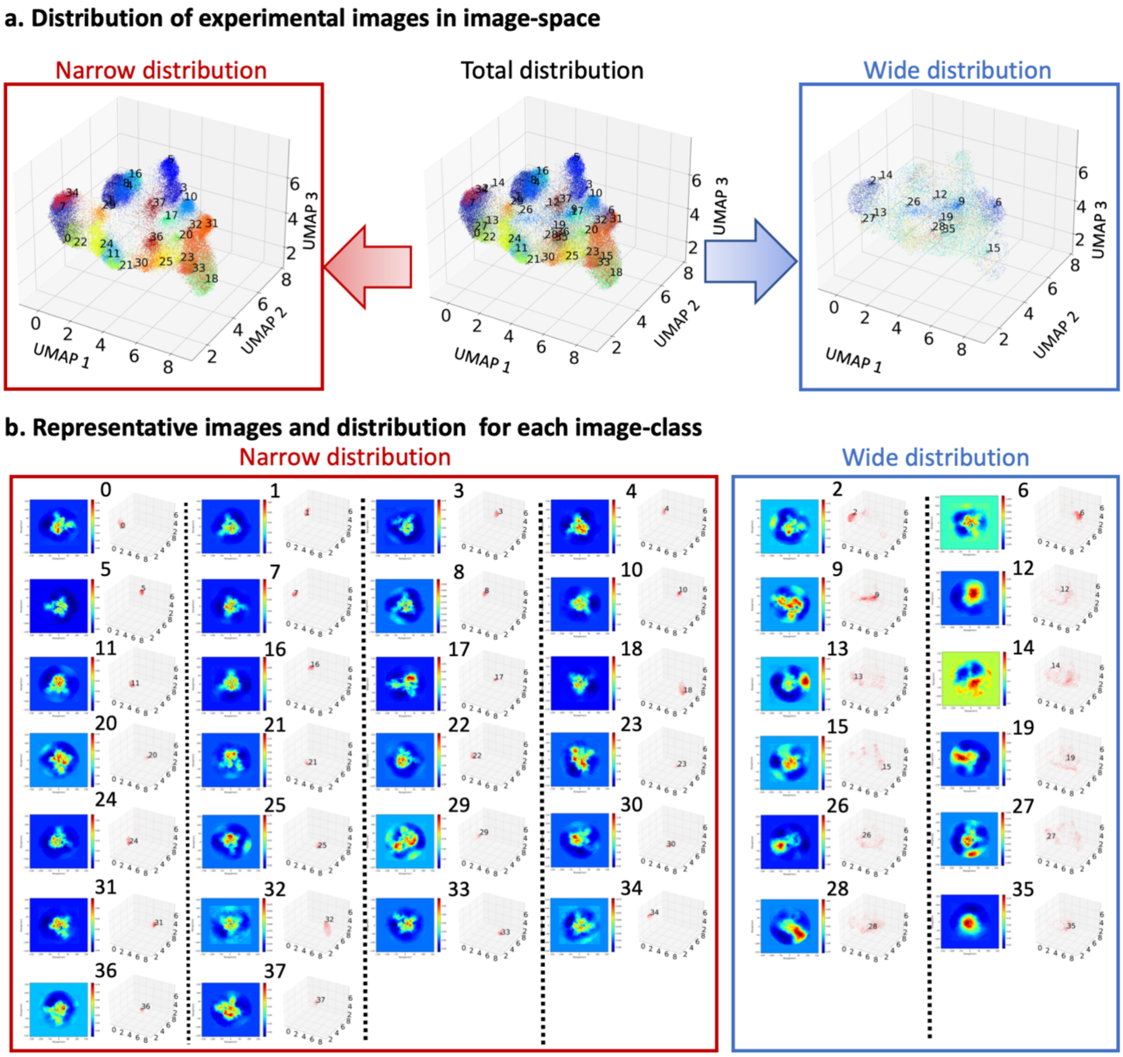


Figure SI3. Image distribution in a 12-dimensional latent space. Displayed in 3D using UMAP. Representative points of VAE-generated images in each cluster are shown. There were clusters of images with narrow distributions and clusters with wide distributions. The clusters with wide distributions tend to include junk images, so in the single-image analysis, we focused on the group of images belonging to the narrow distribution. The representative images, which have the highest probability from GMM clustering, are marked with an "X". Due to memory constraints, approximately ~95,000 points from only the odd-numbered entries were used for the rendering.

**SI4: Results of globule-search for all experimental target image**

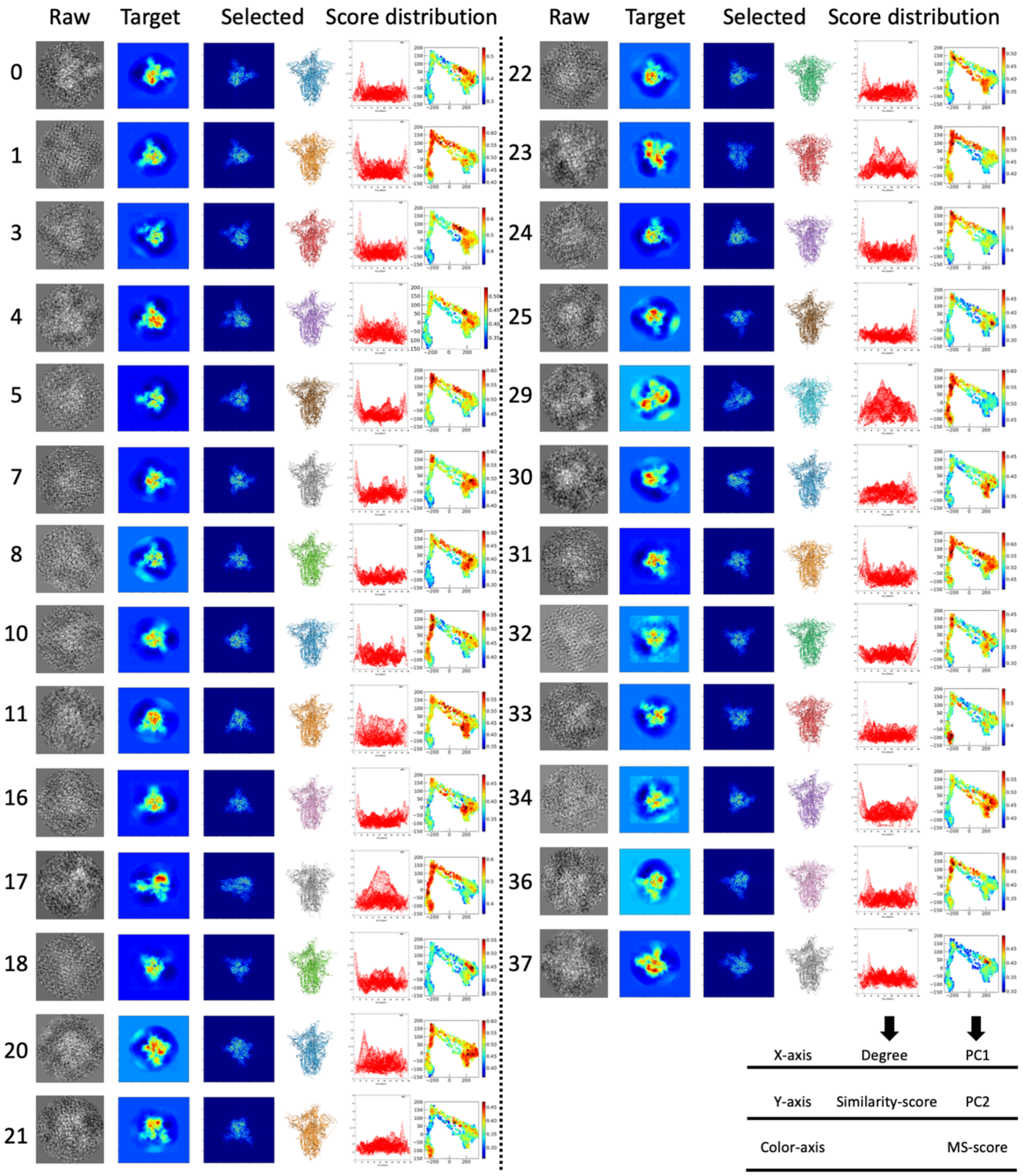


Figure SI4. Results of globule-search for all experimental target image. Structural search was conducted for the 2,580 structures prepared for global structural search. Raw: cryoEM experimental images. Target: target images improved by VAE. Selected: selected 2D images and 3D structures from the search results. Score distribution: (left) Results of molecular orientation search when the MS-score is maximized. X-axis: Degree, Y-axis: Similarity score; (middle) MS-score distribution. X-axis: PC1, Y-axis: PC2; (right) RMSD dependence of MS-score. X-axis: RMSD, Y-axis: MS-score.

**SI5: Four typical structural states estimated by the cryoMDM method**

As examples of four typical structural states, figure SI5 details the results of local structural search targeting T33, T5, T0, and T25. Each column presents the results of 100 cycles of PaCS-MD conducted on the VAE target images a-d. Observing the score progression shown in i-l (X: RMSD vs. Y: matching score), it is evident that local search is conducted within approximately 3 Å around the structures selected from the global search. Notably, in the local search for T0, the search extends beyond the global search, clearly highlighting the necessity of local search. The structures obtained through local search with MS-scores are shown in m-p. T5 represents a typical Close state, while T25 corresponds to a typical Open state. Additionally, the estimated structure for T33 was in a Condensed state, with the RBD closely interacting with surrounding domains. In contrast, T0 exhibited a Free state for the RBD. These results suggest that the spike protein does not merely oscillate between Open and Close states in a non-binding state, but instead undergoes continuous structural changes, traversing various intermediate states.

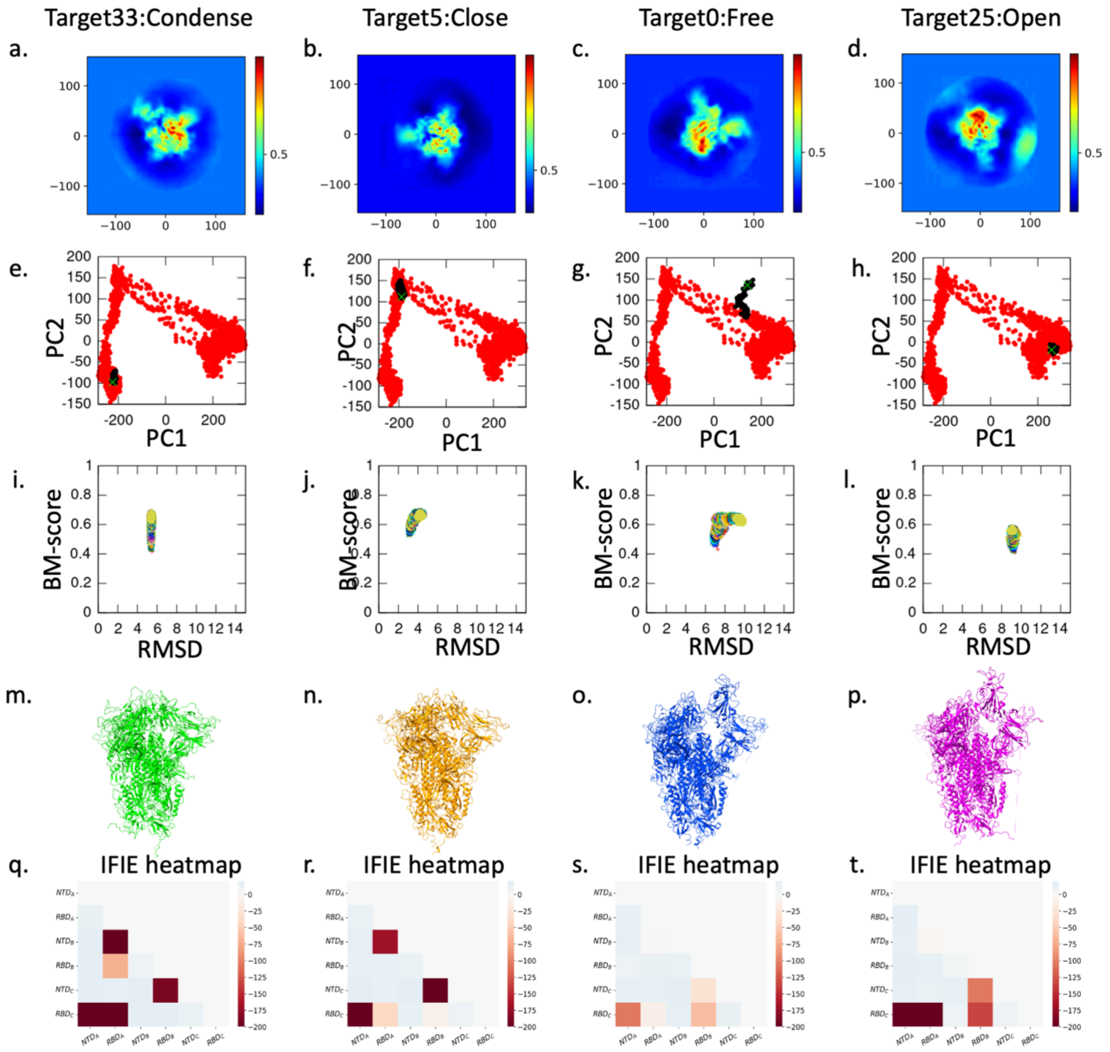

Figure SI5. Results of structural search using target images for T33, T5, T0, and T25. The first row shows the target images (VAE images), the second row presents the distribution of structures from global search (red dots), along with the PaCS-MD progression (black dots) and the location of the highest score structure (marked with an X). The third row illustrates the score progression (X: RMSD vs. Y: matching score). The fourth row displays the all-atom structure with the highest score of Condense, Close ,Free and Open states. The fifth row features the IFIE heatmap for each domain.

**SI6: Results of FMO calculations for the four typical structural states**

The diverse 3D atomic model structures obtained by the cryoMDM method allow for detailed structural evaluation. Here, quantum chemical calculations were performed using the Fragment Molecular Orbital (FMO) method to precisely assess the intramolecular interactions at the epitope site. To stably evaluate the estimated structures through the FMO method, ensemble FMO calculations were conducted using the Top 10 structures from the aforementioned local search. FMO calculations were performed using ABINIT-MP for a total of 260 structures, combining 26 estimated structures and the Top 10 structures. FMO calculations converged for a total of 245 structures, and an Interfragment Interaction Energy (IFIE) analysis was conducted to obtain the ensemble average of IFIE for the 26 estimated structures (See Method).

The spike protein forms a trimer, and we focused on the S1 region, which contains the epitope where ACE binds. We defined structural domains by subdividing the S1 region into [NTD: 1-302], [RBD: 333-527], and [SD: 532-685], resulting in domains ($RBD_A$, $RBD_B$, $RBD_C$, $NTD_A$, $NTD_B$, $NTD_C$) and conducted interaction analyses between these domains. The results of the Interfragment Interaction Energy (IFIE) analysis for the 26 estimated structures are shown in Figure 4. The inter-domain IFIE map is composed of a 6×6 tile arrangement, where the redder the color of each tile, the stronger the stable interaction between the two domains. The domains that indicate important interactions in structural changes are $RBD_A$, $RBD_B$, $RBD_C$, and $NTD_B$, with four distinct IFIE patterns observed.

As representative examples of each state, the interfragment interaction energy (IFIE) maps for the aforementioned clusters 5, 25, 0, and 33 are shown in Figure SI6 a-b. The estimated structure of the target image for Cluster 5 displays a typical IFIE pattern in the Close state (Figure SI6 b), indicating that in the Close structure, $RBD_A$ is stabilized through interaction with $NTD_B$. Similarly, $RBD_B$ and $RBD_C$ are stabilized through interactions with $NTD_C$ and $NTD_A$, respectively. In contrast, examining the typical IFIE pattern in the Open state represented by the estimated structure of Cluster 25 (Figure SI6 d) reveals that the stabilization between the $RBD_A$ and $NTD_B$ domains is lost, and a stable interaction occurs between $RBD_A$ and $RBD_C$. Additionally, stable interactions also arise between $RBD_C$ and $RBD_B$ during this state. Furthermore, the structures estimated from Cluster 0, which is positioned at the transition between Open and Close structures, show very weak inter-domain interactions as illustrated in Figure SI6c. This suggests that in the intermediate state during the transition between Open and Close structures, the RBD is in a highly free state. Moreover, Cluster 33, as shown in Figure SI6a, exhibits both types of inter-domain interactions characteristic of the aforementioned Close and Open structures, indicating that it is in a condensed state.

To analyze the interactions of the representative structures of the four characteristic structural states (clusters 33, 25, 0, and 5), a PIEDA analysis was performed (Figure SI6). Among the energy components obtained from the PIEDA analysis (ES, EX, CT+mix, DI), pairs of fragments were defined as important fragments if the absolute value of any of the three components (short-range interactions) other than ES was 3 kcal/mol or greater. Important fragments for the representative structures of the four structural groups (clusters 33, 25, 0, and 5) are highlighted in yellow in Figures SI6e-p. The important fragments differ across the four characteristic structural states, indicating that the intramolecular interactions are continuously transitioning to form quasi-stable intermediate structures.

Based on the occurrence frequency of important fragment pairs, we identified the key fragments that characterize each state. The fragments that interact most frequently and contribute to structural stabilization among the various structural domains were compiled into the Most Frequent Fragment List in Table SI1. For each inter-domain interaction, the most frequently occurring fragments (residues) for each structural domain are listed in the corresponding rows. This approach successfully demonstrated the intramolecular interactions that significantly contribute to the structural stabilization of the four characteristic structural states at the residue level, grounded in CryoEM individual experimental data. This information is expected to provide useful insights for designing inhibitors in structure-based drug design (SBDD).

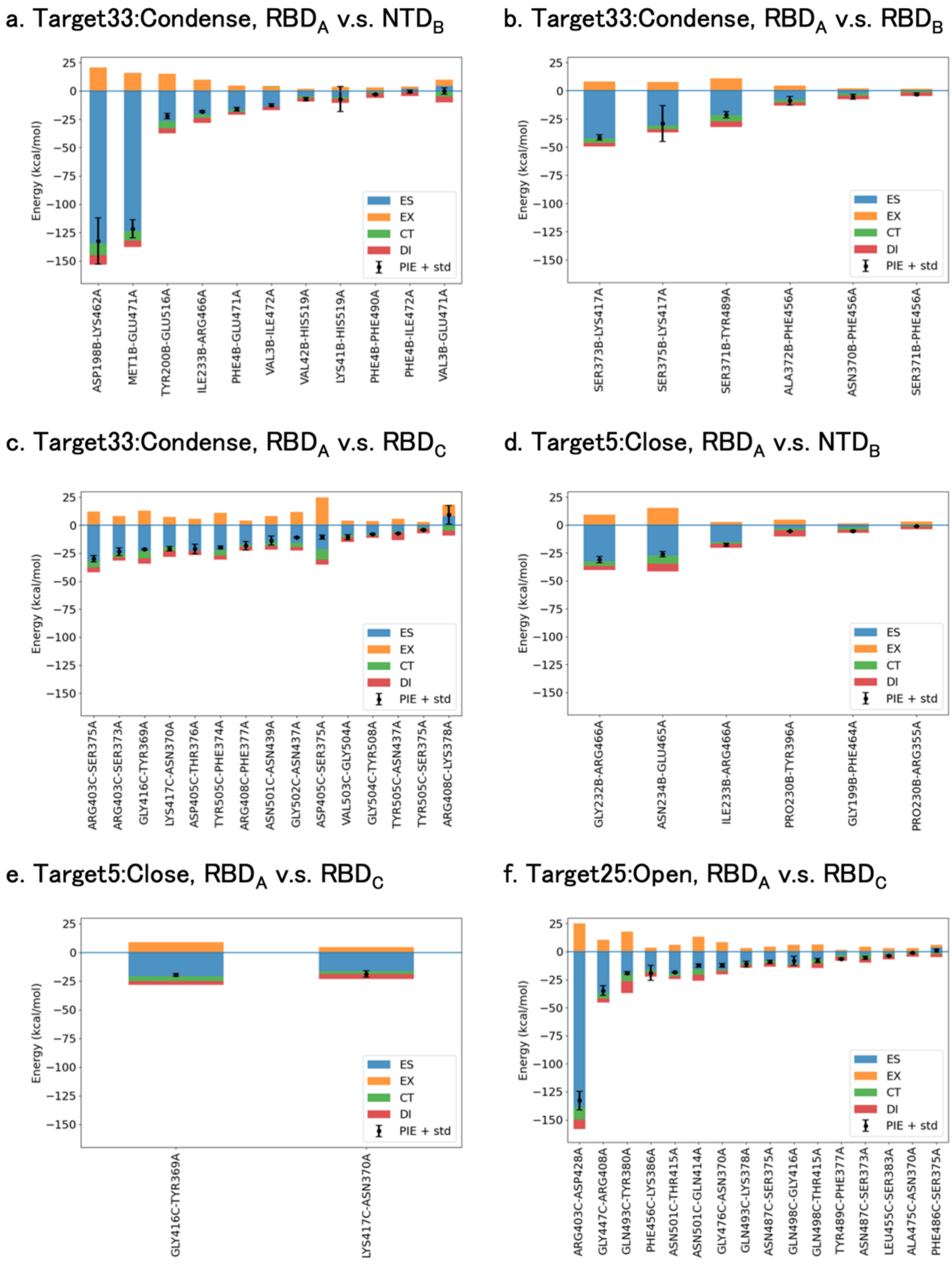


Figure SI6. Results of FMO calculations for the four typical structural states of Condense, Close, Free, and Open states (Targets 33,5,0 and 25) estimated by the cryoMDM method. a-d. IFIE heatmap diagrams. e-q. Important interaction fragments.

**SI7: The RMSD plots of the top 10 scored structures estimated from each target image using cryoMDM methods**

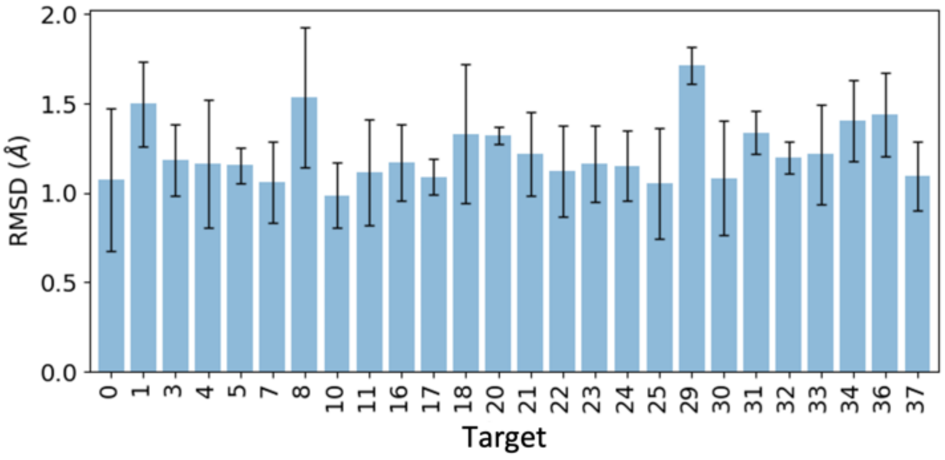


Figure SI7. The structures with the highest score for each Target was used as the reference structure, and the RMSD was calculated for the remaining nine structures. The average and standard deviation of the RMSD for each Target are presented. X-axis: Target image ID, Y-axis: RMSD.

### SI8: Results of PIEDA analysis

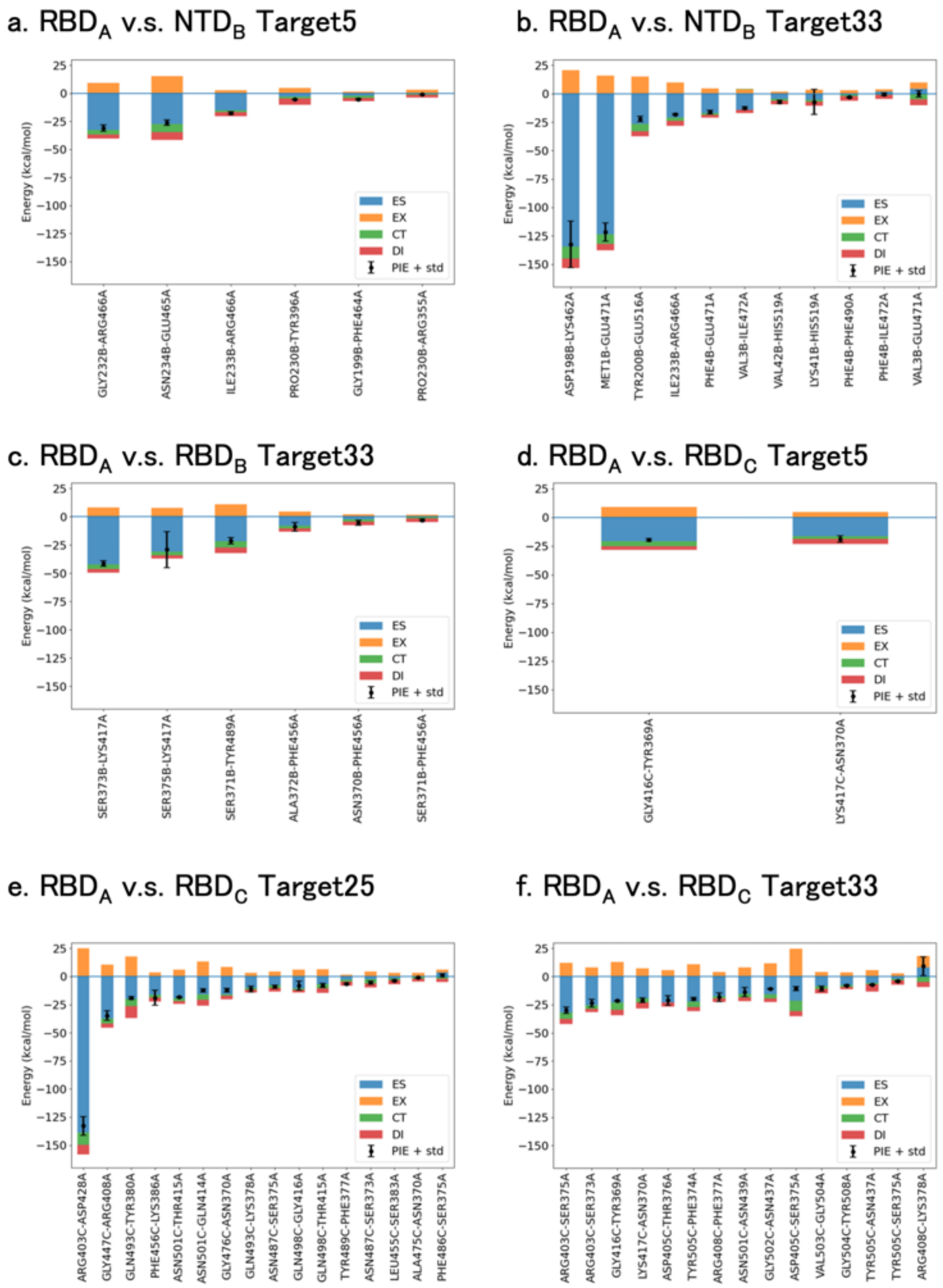


Figure SI8. Results of PIEDA analysis for the four typical structural states of Condense, Close, Free, and Open states (Targets 33,5,0 and 25). The free state of Target0 has no figure because there were no important residues.

---

**SI9: List of important residues obtained from PIEDA analysis**

For interactions between domains I and J, the most frequently occurring residues in each domain are listed in rows corresponding to the respective domains. If multiple residues are mentioned within the same cell, it indicates they occurred with the same frequency. "n/a" denotes that there are no close interactions in the respective domain pair. The numbers in parentheses represent the occurrence frequency. (a) shows results for the Condense state (Targets: 17, 23, 31, **33**), (b) for the Close state (Targets: **5**, 10, 22, 24, 29, 32, 36), (c) for the Free state (Targets: **0**, 1, 3, 4, 21, 37), and (d) for the Open state (Targets: 7, 8, 11, 16, 18, 20, **25**, 30, 34).

The cells in the table are defined by the following rules:

1. Cells (Row A, Column B) and (Row B, Column A) both represent important residues involved in close interactions between domains A and B.
2. Cell (Row A, Column B) contains the residue with the highest occurrence frequency among the important residues belonging to domain A. The same applies to cell (Row B, Column A).
3. The numbers in parentheses indicate the number of times each residue has occurred.

(a) The condense state（Target: 17, 23, 31, **33**）

| | $NTD_A$ | $RBD_A$ | $NTD_B$ | $RBD_B$ | $NTD_C$ | $RBD_C$ |
|---|---|---|---|---|---|---|
| $NTD_A$ | | n/a | n/a | n/a | n/a | ASN234A<br>(5) |
| $RBD_A$ | n/a | | LYS462A<br>GLU471A<br>(5) | LYS417A<br>(6) | n/a | TYR369A<br>SER375A<br>(4) |
| $NTD_B$ | n/a | TYR200B<br>(4) | | n/a | n/a | n/a |
| $RBD_B$ | n/a | ASN370B<br>(4) | n/a | | GLU516B<br>ARG466B<br>(4) | LYS417B<br>(3) |
| $NTD_C$ | n/a | n/a | n/a | LYS41C<br>ILE233C<br>ASN234C<br>(3) | | n/a |
| $RBD_C$ | TYR396C<br>(5) | LYS417C<br>(5) | n/a | ASN370C<br>(2) | n/a | |

(b) The close state (Target: **5**, 10, 22, 24, 29, 32, 36)

| | $NTD_A$ | $RBD_A$ | $NTD_B$ | $RBD_B$ | $NTD_C$ | $RBD_C$ |
|---|---|---|---|---|---|---|
| $NTD_A$ | | n/a | n/a | n/a | n/a | ILE233A (9) |
| $RBD_A$ | n/a | | ARG466A (8) | LYS417A (7) | n/a | ASN370A (6) |
| $NTD_B$ | n/a | PRO230B (7) | | n/a | n/a | n/a |
| $RBD_B$ | n/a | ASN370B (9) | n/a | | GLU465B (7) | LYS417B (5) |
| $NTD_C$ | n/a | n/a | n/a | ASN234C (7) | | n/a |
| $RBD_C$ | ARG466C (12) | LYS417C (5) | n/a | ASN370C (3) | n/a | |

(c) The free state (Target: **0**, 1, 3, 4, 21,37)

| | $NTD_A$ | $RBD_A$ | $NTD_B$ | $RBD_B$ | $NTD_C$ | $RBD_C$ |
|---|---|---|---|---|---|---|
| $NTD_A$ | | n/a | n/a | n/a | n/a | TYR200A (10) |
| $RBD_A$ | n/a | | ARG357A (2) | ASN460A (2) | n/a | LYS378A (7) |
| $NTD_B$ | n/a | TYR200B GLY232B (2) | | n/a | n/a | n/a |
| $RBD_B$ | n/a | SER371B ALA372B SER373B SER375B LYS378B (1) | n/a | | ARG357B (5) | LYS417B (10) |
| $NTD_C$ | n/a | n/a | n/a | TYR200C (4) | | n/a |
| $RBD_C$ | ARG357C (7) | PHE486C (9) | n/a | TYR369C ASN370C THR376C (4) | n/a | |

(d) The open state（Target: 7, 8, 11, 16, 18, 20, **25**, 30, 34）

| | NTD$_A$ | RBD$_A$ | NTD$_B$ | RBD$_B$ | NTD$_C$ | RBD$_C$ |
|---|---|---|---|---|---|---|
| NTD$_A$ | | n/a | n/a | n/a | n/a | TYR200A (4) |
| RBD$_A$ | n/a | | ARG357A (7) | n/a | n/a | LYS386A (11) |
| NTD$_B$ | n/a | PHE168B (6) | | n/a | n/a | n/a |
| RBD$_B$ | n/a | n/a | n/a | | ARG357B (13) | ASP405B (13) |
| NTD$_C$ | n/a | n/a | n/a | TYR200C (14) | | n/a |
| RBD$_C$ | GLU516C (4) | (10) | n/a | LYS378C (13) | n/a | |

**SI10: Conducting of cryoDRGN analysis**

Using the J60 dataset (19,3125 images) applied in the cryoMDM method for spike protein, we conducted a CryoDRGN analysis. The hyperparameters for CryoDRGN were set as follows: $\beta$ = 0.8, latent space dimension: zdim = 12, batch size = 64, learning rate: Ir = 0.0001, and number of epochs = 50. The empirical distribution in the latent space is shown in Figure 5c using UMAP. Additionally, density maps for 20 representative points are displayed. In the examples illustrated in Figure 5c, all density maps indicate an open state, suggesting that polymorphism is less likely to be produced in this case.

**SI11: Experimental movie**

(/vol0006/hp200178/data/20210930_OsakaU_VAE_analysis/Matching_OsakaU_J60/cluster_kiridashi_multiTarget_Analysis2_OsakaU_J60_180pixel_cluster18）

clusetr0:TMD3_toUp

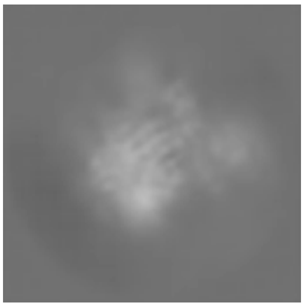

・cluster5:MD2_Down

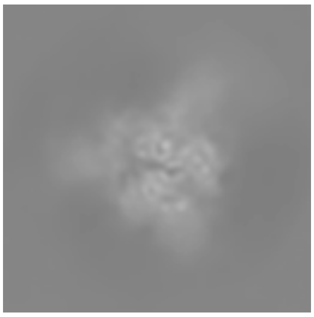

・cluster25:MD1_Up

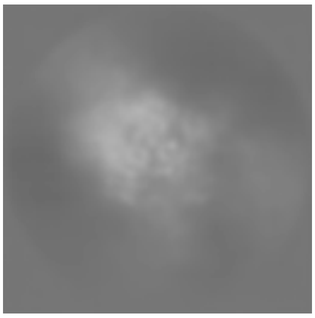

・clusetr33:MD1_Down

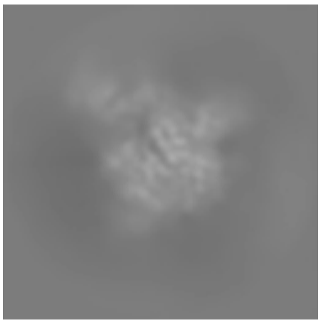

**SI12: VAE Network Model**

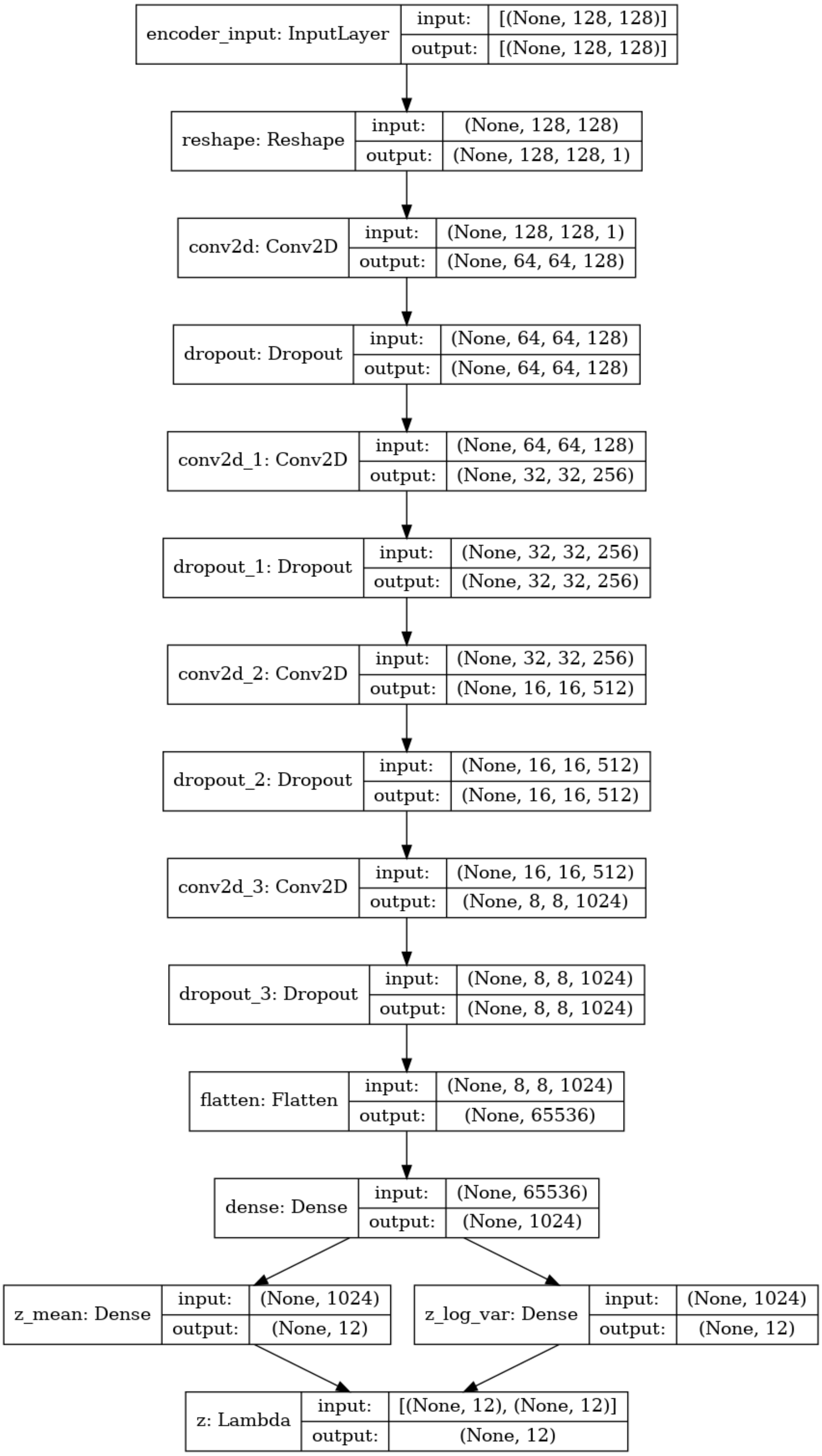

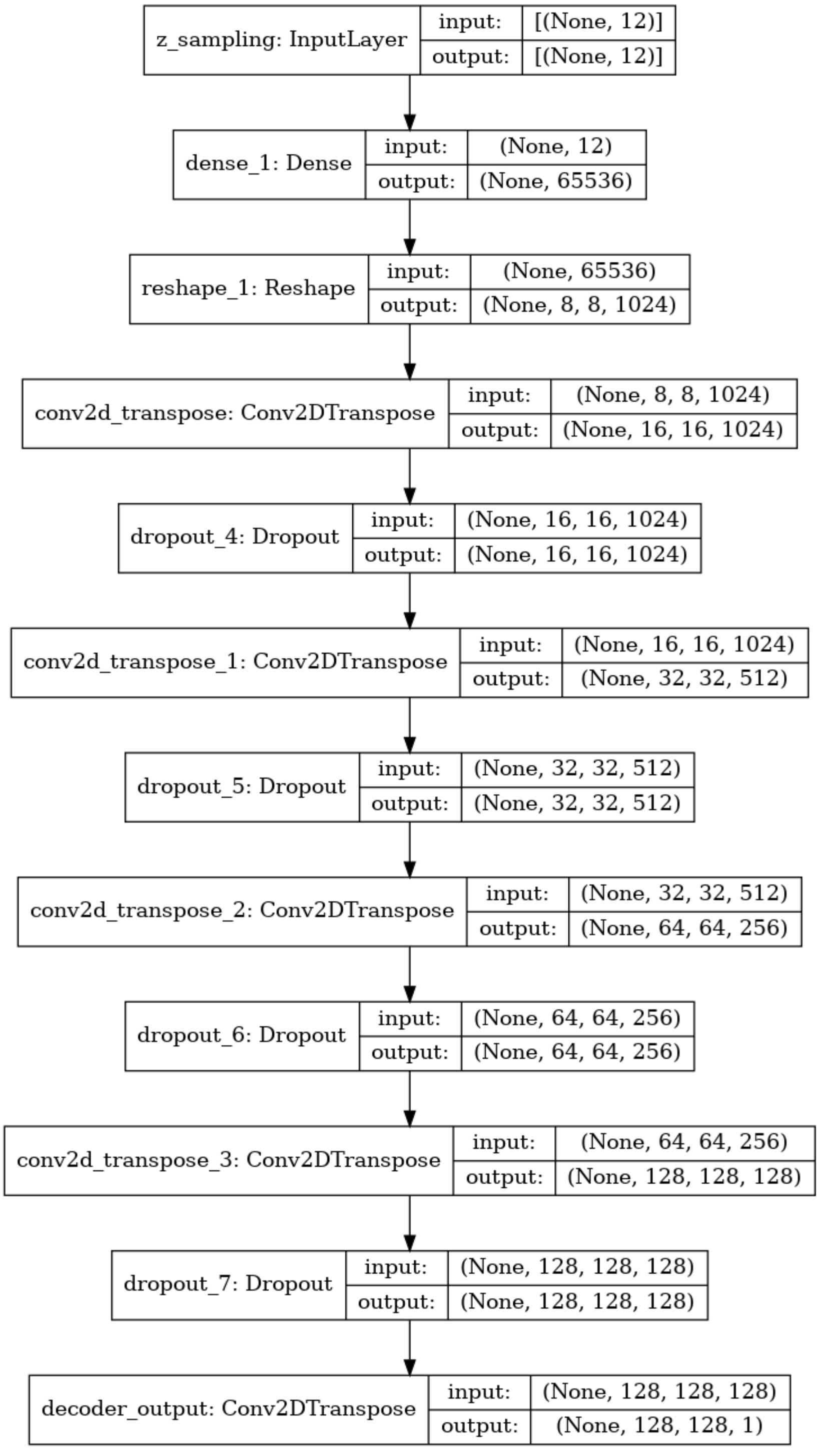

z_sampling: InputLayer
input: [(None, 12)]
output: [(None, 12)]
dense_1: Dense
input: (None, 12)
output: (None, 65536)
reshape_1: Reshape
input: (None, 65536)
output: (None, 8, 8, 1024)
conv2d_transpose: Conv2DTranspose
input: (None, 8, 8, 1024)
output: (None, 16, 16, 1024)
dropout_4: Dropout
input: (None, 16, 16, 1024)
output: (None, 16, 16, 1024)
conv2d_transpose_1: Conv2DTranspose
input: (None, 16, 16, 1024)
output: (None, 32, 32, 512)
dropout_5: Dropout
input: (None, 32, 32, 512)
output: (None, 32, 32, 512)
conv2d_transpose_2: Conv2DTranspose
input: (None, 32, 32, 512)
output: (None, 64, 64, 256)
dropout_6: Dropout
input: (None, 64, 64, 256)
output: (None, 64, 64, 256)
conv2d_transpose_3: Conv2DTranspose
input: (None, 64, 64, 256)
output: (None, 128, 128, 128)
dropout_7: Dropout
input: (None, 128, 128, 128)
output: (None, 128, 128, 128)
decoder_output: Conv2DTranspose
input: (None, 128, 128, 128)
output: (None, 128, 128, 1)
